# Nonlinear Relationships of Fibrin Network Structure as a Function of Fibrinogen and Thrombin Concentrations for Purified Fibrinogen and Plasma Clots

**DOI:** 10.64898/2026.01.22.700990

**Authors:** Can Cai, Zezhong Zhang, Arezoo Nameny, Daniel Nelli, Keith Bonin, Brittany E. Bannish, Nathan E. Hudson, Glen Marrs, Stephen R. Baker, Martin Guthold

**Affiliations:** Department of Physics, Wake Forest University, Winston-Salem, NC 27109; Department of Mathematics and Statistics, University of Central Oklahoma, Edmond, Oklahoma, 73034; Department of Physics, East Carolina University, Greenville, NC 27885; Department of Biology, Wake Forest University, Winston-Salem, NC 27109

**Author notes:** These authors are co-corresponding authors of this article. These authors contributed equally to this work (Can Cai, Zezhong Zhang).

**Keywords:** Biopolymer Network Structure, Fibrin Network Structure, Fibrinogen, Thrombin

## Abstract

**Background:** Fibrinogen levels are associated with bleeding disorders and thrombotic disease. Thrombin converts fibrinogen to fibrin, producing the load-bearing fibrin scaffold that governs clot mechanics and transport.

**Objective:** Quantitatively map how initial fibrinogen and thrombin concentrations, [*Fgn*]_0_ and [*T*ℎ*r*]_0_, determine fibrin architecture in human plasma and a purified fibrinogen system.

**Methods:** Scanning electron microscopy was used to quantify single-fiber morphology—fiber diameter and branch-to-branch segment length from a standardized sample-preparation protocol. Confocal microscopy was used to quantify network architecture—projected fiber density and pore/bubble size.

**Results and Conclusions:** Across plasma and purified systems, fibrin structural parameters were quantitatively captured by compact multiplicative scaling laws of the form *Y* = *k*[*Fgn*]_0_*^α^*[*T*ℎ*r*]_0_*^β^*. Unlike prior studies, which examined narrower condition ranges without establishing predictive equations across a systematic fibrinogen–thrombin concentration matrix, this framework defines distinct, quantitative roles for fibrinogen and thrombin in fibrin assembly. The magnitudes and signs of the exponents α and β indicate that thrombin primarily controls individual fiber growth kinetics, strongly shortening branch-to-branch segment length and modestly thinning fibers, whereas fibrinogen primarily controls space filling, strongly increasing fiber density, reducing pore/bubble size, and thickening fibers. For matched [*Fgn*]_0_ and [*T*ℎ*r*]_0_, compared to plasma clots, purified fibrinogen formed denser networks with thinner and shorter fibers, suggesting that the plasma biochemical environment partially inhibits thrombin activity. Fiber length analysis further suggests that each thrombin molecule nucleates one fiber segment. Together, these parameterized scaling relations provide a predictive quantitative framework linking clot composition to fibrin microstructure in plasma and purified fibrinogen clots.

## Introduction

Blood clotting is tightly regulated in space and time: clots must assemble rapidly and locally at sites of vascular injury to prevent blood loss, seal the wound, and limit infection. In bleeding disorders such as hemophilia and von Willebrand disease, impaired fibrin formation or platelet adhesion can yield delayed, undersized, mechanically weak clots that may fail to arrest bleeding. When clotting is initiated inappropriately or fails to undergo timely fibrinolysis, persistent thrombi in critical vascular beds such as the brain, heart, and deep veins can lead to ischemic stroke, myocardial infarction, and deep vein thrombosis [1, 2].

Fibrin fibers provide the load bearing scaffold of the clot. In the prevailing assembly model, thrombin cleaves fibrinopeptides A and B on fibrinogen to expose binding sites termed ‘knob A’ and ‘knob B’ that drive knob-hole interactions, forming fibrin aggregates and half staggered protofibrils that elongate by end addition and thicken by lateral aggregation, a process influenced by the fibrin αC regions. As these bundles intersect, they form a branched, three-dimensional network. Abnormal fibrinogen levels are clinically relevant: elevated fibrinogen is associated with ischemic stroke, myocardial infarction, and venous thromboembolism, whereas reduced fibrinogen, for example in postpartum hemorrhage, creates a predisposition to bleeding [3–5]. Thrombin generation is likewise central: insufficient thrombin, as in some types of hemophilia, causes hemostatic failure, while excessive or sustained thrombin, as in cancer associated thrombosis or antiphospholipid syndrome, promotes thrombosis [6–8]. Together, fibrinogen and thrombin set the structural state of the network, which in turn shapes clot mechanical behavior and susceptibility to lysis [1].

A persistent challenge is that published structure-concentration relationships are difficult to compare across studies, even when the qualitative conclusions appear similar. For fiber diameter, most reports spanning confocal microscopy, SEM, and turbidimetry support an increase in fiber diameter with increasing fibrinogen, whereas a smaller set report a decrease in fiber diameter with increasing fibrinogen [9–13]. Reported diameters also span a wide range, from tens of nanometers to micrometers [10–13]. This variability is not solely biological; it reflects differences in measurement resolution and in the analysis and modeling frameworks used to map protein concentration to structure. Confocal microscopy is diffraction limited and cannot resolve fiber diameters under about 300 nm [14]. SEM provides higher spatial resolution, but sample preparation steps such as washing, dehydration, drying and metal coating can alter apparent geometry [9, 11–13]. Light scattering relies on equations that use restrictive approximations and can be less accurate when fibers are very thin or become very thick (on the order of the wavelength of light) or networks are highly heterogeneous [10]. As a result, studies have described diameter dependence using different functional forms, including approximately linear relations in some SEM analyses and nonlinear scaling consistent with power law behavior in scattering-based work [10, 12]. In contrast to the fibrinogen dependence, there is broader agreement that increasing thrombin produces thinner fibers, although the reported magnitude again varies across methods [9, 11–13].

Fiber length and pore scale metrics add further complexity because they are intrinsically harder to define and measure in images. Fiber segments commonly extend beyond the field of view, and apparent crossings in two-dimensional projections do not always correspond to true branch points, complicating segmentation and length assignment [15]. A prior confocal study quantified apparent lengths from single planes and reported shorter apparent lengths with increasing fibrinogen or thrombin [12]. However, a growing body of work indicates that structural features such as fiber diameter and length are not well described by normal distributions, motivating robust summary statistics and careful aggregation across images [9, 16, 17]. For example, a standardized SEM study reported lognormal diameter distributions [17], and three-dimensional confocal reconstructions indicate lognormal-like behavior for both diameter and length [9]. For network packing, fiber density has been estimated from confocal images using intensity-based measures [12] and from light scattering through inference of structural parameters [10]. Pore size is often quantified from confocal data using a largest inscribed circle or bubble analysis that yields a pore diameter distribution [18], alongside alternative estimators such as chord length intercepts or distance map-based approaches [19]. Across these approaches, a consistent qualitative trend is that higher fibrinogen or thrombin tends to reduce pore size, although the reported functional form varies between datasets and analysis choices [12, 18, 20, 21].

Despite extensive prior work describing how fibrinogen and thrombin influence fibrin network architecture, the field still lacks a predictive and quantitatively transferable mapping from composition to structure that spans multiple structural metrics and can be applied across studies. In practice, most reports emphasize qualitative trends (e.g., thicker versus thinner fibers) or provide method-specific absolute values that are difficult to reconcile because definitions, resolution limits, and preparation/analysis workflows differ. This limits mechanistic inference and modeling: without internally consistent scaling relationships for fiber diameter, segment length, network density, and pore size, it remains difficult to (i) test assembly hypotheses, (ii) link microstructure to mechanics and fibrinolysis, or (iii) calibrate and validate multiscale simulations of clot structure, mechanics, and transport.

Here, we test the hypothesis that fibrin architecture across a systematic fibrinogen–thrombin concentration matrix follows a common nonlinear scaling form that can be compactly expressed as power-law relationships of the form *k*[*Fgn*]_0_*^α^*[*T*ℎ*r*]_0_*^β^*. We investigate both plasma and purified fibrinogen systems, enabling direct cross-system comparisons. We combine standardized high-magnification SEM [17] to quantify fiber diameter and branch-to-branch segment length with 3D confocal imaging to quantify fiber density and pore size [18]. Using a shared power-law parameterization across metrics and systems, we derive a unified set of scaling equations that serves as a practical “structure calculator” for predicting fibrin network morphology and provides quantitative benchmarks for mechanistic assembly models and computational simulations of fibrin and related fibrous hydrogels. Unlike prior studies that examined more limited conditions, this work establishes predictive equations across a systematic fibrinogen–thrombin concentration matrix and reveals distinct quantitative roles of fibrinogen and thrombin in fibrin assembly. Similar exponents across the two systems for all measured quantities suggest a common underlying assembly mechanism, whereas differences in the scaling prefactor *k* are plausibly attributable to reduced effective thrombin activity in plasma. Exponent magnitudes indicate effect size: increasing fibrinogen primarily increases fiber density, decreases pore size, and thickens fibers, with little effect on segment length, whereas increasing thrombin increases fiber density, decreases pore size, and produces thinner, shorter fibers. The combined measurements of fiber density and single-fiber volume allow estimation of the fibrin-occupied volume fraction within the total sample volume and suggest that thicker fibers may have lower mass density. Finally, segment length analysis suggests that each thrombin molecule may nucleate a single fiber segment.

## Materials & Methods

### Reagents

Pooled normal human plasma (George King Bio-Medical, Product #0010) and Peak 1 (purified fibrinogen) human fibrinogen (Enzyme Research Laboratories, Cat# P1 FIB) were aliquoted and stored at −80 °C. Human thrombin (Enzyme Research Laboratories, Cat# HT1002a; lot-specific activity: 3044 NIH U/mg) was aliquoted at 200 NIH U/ml and stored at −80 °C. Alexa Fluor 488–conjugated human fibrinogen (Thermo Fisher Scientific, Cat# F13191) was aliquoted and stored at −80 °C. Aliquots were thawed at 37 °C immediately before use.

Thrombin concentrations are reported in NIH U/mL because the stock solution was supplied and diluted according to calibrated activity units; the molar concentration is provided only as a lot-specific approximate conversion.

### SEM sample preparation and imaging

#### Clot conditions

Plasma clots were prepared at room temperature across a 4×4 matrix of fibrinogen (0.36, 0.73, 1.45, 2.9 mg/mL) and thrombin (0.05, 0.1, 0.2, 1.0 U/mL) concentrations. Purified fibrinogen clots were prepared across a 3×3 matrix of fibrinogen (0.36, 0.73, 1.45 mg/mL) and thrombin (0.05, 0.1, 0.2 U/mL) concentrations. The plasma 4×4 matrix was reduced to 3×3 for purified clots because clots formed at the highest fibrinogen and thrombin concentrations were too dense to quantify reliably. Clotting was initiated by adding an activation mix containing thrombin and CaCl₂ to achieve the thrombin concentrations listed above in Tris-buffered saline (50 mM Tris, 100 mM NaCl, pH 7.4). CaCl₂ was included at 20 mM for plasma clots and 9.5 mM for purified fibrinogen clots. The plasma was supplied as 3.2% (w/v) sodium citrate–anticoagulated plasma (corresponding to ∼11 mM citrate in the final plasma). The higher CaCl₂ in plasma was used to recalcify citrated plasma (George King) and overcome citrate chelation, whereas purified fibrinogen contains no citrate; we therefore used 9.5 mM CaCl₂ in the purified system to keep the added Ca²⁺ in the same order of magnitude and facilitate comparison between plasma and purified conditions. Clots were allowed to polymerize for 2 h at room temperature.

#### Fixation, drying, and coating

Samples were prepared using a standardized wash–fix– dehydrate–dry workflow as previously described [17]. After polymerization, clots formed in Eppendorf tube lids were washed three times in sodium cacodylate buffer (50 mM sodium cacodylate, 100 mM NaCl) and then fixed by immersion in 2 mL of 2% glutaraldehyde prepared in the same buffer for 6 h. Following fixation, samples were washed three additional times in sodium cacodylate buffer, dehydrated through a graded ethanol series (30%-100%), and dried using hexamethyldisilazane (HMDS). Dried samples were sputter-coated with gold for 45 s (Cressington Model 108).

#### SEM imaging

Samples were imaged on a Zeiss Gemini 300 SEM under high vacuum at 30,000× magnification. For each condition, two independent clot preparations (biological replicates) were performed. From each replicate, eight micrographs were acquired, yielding 16 images per condition in total. Fields of view were sampled from widely separated, non-overlapping locations distributed across each clot (at fixed 30,000× magnification), while avoiding the film region, as described in [17].

### Confocal sample preparation and imaging

#### Clot conditions and sample preparation

For confocal microscopy, plasma clots were prepared using a 4×4 matrix of fibrinogen (0.36, 0.73, 1.45, and 2.9 mg/mL) and thrombin (0.05, 0.1, 0.2, and 1.0 U/mL) in TBS (20 mM Tris, 150 mM NaCl, pH 7.4). Purified fibrinogen clots were prepared using a 3×3 matrix of fibrinogen (0.36, 0.73, and 1.45 mg/mL) and thrombin (0.05, 0.1, and 0.2 U/mL) in the same buffer. The purified clots were reduced to 3x3 matrix because clots formed at the highest fibrinogen and thrombin concentrations were too dense to quantify reliably. CaCl₂ was included at 20 mM for plasma clots and 9.5 mM for purified fibrinogen clots, as described in the SEM section above. Alexa Fluor 488–fibrinogen was incorporated at 6.5% of the total fibrinogen concentration. Immediately before use, the labeled fibrinogen was warmed and briefly centrifuged (10 s) to remove aggregates. Reactions were loaded into channel slides (ibidi μ-Slide VI 0.4), shielded from light, and allowed to polymerize for 2 h at room temperature.

#### Confocal imaging

Imaging was carried out on a Zeiss LSM 710 laser-scanning confocal microscope with a 40× oil-immersion, 1.3 NA objective lens using 488 nm excitation light, at room temperature. For each condition, 36 z-stacks were collected (18 per slide from two slides). Each slide contained six channels; three imaging positions per channel were selected at random, spaced several millimeters apart and positioned 3–6 μm away from channel edges to reduce boundary effects. Stacks spanned 212 μm × 212 μm in x–y and 15 μm in z (33 optical sections, 0.47 μm step size) and were acquired at 8-bit gray scale. Detector gain was tuned to clearly resolve fibers while minimizing background signal.

Plasma clots with no fluorescently labeled fibrinogen added were imaged under the same 488 nm excitation and detection settings used for the Alexa Fluor 488-fibrinogen samples. No fibrin-like network fluorescence was detected under these conditions.

### Image analysis and quantitative network metrics

#### Fiber diameter

An ImageJ-based manual approach was used to quantify fiber diameter. Briefly, an evenly spaced grid was overlaid on each image, and the fiber intersecting (or closest to) each grid intersection was selected for measurement, ensuring that each selected fiber was measured only once. At least 70 fibers were measured per image, as described in [17].

#### Fiber length

Using the grid-sampling method described above, fiber length was traced in ImageJ and defined as the distance along a fiber segment between adjacent branch points. To avoid including truncated segments near the field boundaries, measurements were taken only within an interior region of interest defined by excluding a 10% margin from each image edge (i.e., fibers were measured only if they lay ≥10% of the image width/height away from the boundaries), while using the original, unaltered images. This corresponds to sampling the central 64% of the field of view. At least 30 fibers were measured per image.

Branch points were identified conservatively based on local fiber morphology in the high-magnification SEM images. Only junctions showing clear continuity, merging, or splitting of fibers within the topmost resolved fiber layer were counted as branch points. Apparent crossings were not automatically counted; crossings in which fibers remained visually separable, appeared to pass over or under one another, were out of focus, or could not be assigned unambiguously to a true junction were excluded. Fiber segments were also excluded if either endpoint was truncated, obscured by fiber pile-up, or located outside the defined interior region of interest.

#### Fiber density

Image stacks were converted to 2D maximum-intensity projections (ImageJ/Fiji) and then binarized using the same thresholding procedure across conditions. To estimate an in-plane fiber density, twenty horizontal and twenty vertical equally spaced line probes were overlaid on each projection, and the number of visible fibrin fibers intersecting each line was counted. Fiber density is initially measured as the mean number of intersections per 100 µm line. Because each maximum projection was generated from a 15 µm z-stack, each 100 µm line integrates fluorescence from an effective 100 µm (length) × 15 µm (depth) sampling region; therefore, densities are reported as intersections per 1500 µm². A 15 µm acquisition depth was chosen as a practical compromise that yielded a high number of clearly resolvable, predominantly straight fiber features with minimal background.

#### Bubble analysis

Individual z-slices were exported as multi-page TIFFs and binarized in MATLAB using a threshold set to three times the mean slice intensity. Fiber-free pores were quantified with a custom algorithm that places the largest possible circles (“bubbles”) within void regions on each slice, as described in Molteni et al [18, 22]. Circles that touched fibers, overlapped with larger accepted circles, or intersected the image boundary were discarded. The diameters of the remaining circles were recorded as a proxy for network porosity, where larger bubble diameters indicate larger pore/void regions.

#### Curve fitting and statistical analysis

For each experimental condition, image-level measurements were summarized and related to the two experimental factors, initial fibrinogen concentration [*Fgn*]_0_ and initial thrombin concentration [*T*ℎ*r*]_0_, using a power-law model:

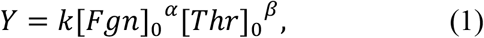

where *Y* denotes the response variable (fiber diameter, fiber length, fiber density, or bubble diameter), k, α, and β are fitted parameters. Model fitting was performed in log space by linear regression of

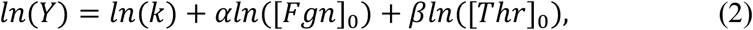

using ordinary least squares (OLS). Goodness of fit was reported as R^2^ computed in log space. For the structural metrics, each image was quantified separately and condition-level summary values were then used for fitting. For confocal analysis of fiber density and bubble diameter, 36 image-level replicates were analyzed per condition. For SEM analysis of fiber diameter and fiber length, 16 image-level replicates were analyzed per condition. Because the distributions of fiber diameter, fiber length, and bubble diameter were non-Gaussian, these responses were summarized for each condition as the median of the image-level medians prior to model fitting. In contrast, fiber density was approximately normally distributed and was summarized for each condition as the mean of the image-level means prior to fitting.

Parameter uncertainty was quantified using bootstrap 95% confidence intervals. Bootstrap resampling was performed at the image level within each condition. In each of 2000 bootstrap iterations, the image-level summary values within each condition were resampled with replacement, a bootstrap condition-level summary value was recalculated, and the full set of condition-level values was refit to the power-law model. The 2.5th and 97.5th percentiles of the resulting bootstrap distributions of *k*, *α*, and *β* were taken as the 95% confidence intervals.

Two-sided p-values were computed from coefficient *t*-tests in the log-space regression for the null hypotheses *H*_0_: ln *k* = 0, *H*_0_: *α* = 0, and *H*_0_: *β* = 0. These tests used the residual degrees of freedom of the corresponding regression model and rely on the usual linear-model assumptions in log space, including independent residuals with approximately constant variance and approximate normality.

For interpretability of the fitting parameters (in the discussion), several model-derived comparisons were calculated from the fitted power-law parameters. Because the model was fit as a two-factor concentration surface, 2^α^ represents the fitted effect of doubling fibrinogen while holding thrombin fixed, and 2^β^ represents the fitted effect of doubling thrombin while holding fibrinogen fixed. The corresponding percent changes were calculated as (2^α^ − 1) × 100% and (2^β^ − 1) × 100%, respectively. Plasma–purified prefactor ratios were calculated by comparing fitted k values for the same structural metric using the same concentration and response-variable units; these ratios represent vertical offsets between the fitted plasma and purified-fibrinogen power-law surfaces.

To assess robustness with respect to the limited number of sampled fibrinogen–thrombin conditions, we performed a leave-one-condition-out analysis(see supplement Table S5 and S6). Each fibrinogen–thrombin condition was omitted in turn, and the power-law fit was repeated using the remaining condition-level summary values. The resulting variation in the fitted parameters was used to evaluate whether the fitted relationships were strongly influenced by any single sample condition.

## Results

### Purified and plasma clots show distinct architectures across matched concentration matrices

Across the fibrinogen–thrombin matrices, SEM and confocal imaging reveal systematic shifts in fibrin network architecture. Under matched conditions, purified clots have thinner, shorter fibers than plasma clots (Figs. 1 and 2; Tables S1 and S2) and form denser, smaller-pore networks (Figs. 3 and 4; Tables S3 and S4). Overall, fibrinogen more strongly modulates fiber density and pore size, whereas thrombin more strongly influences fiber diameter and length, as it can be seen by the magnitude of the exponents in Tables 1 and 2.

**Figure 1:**
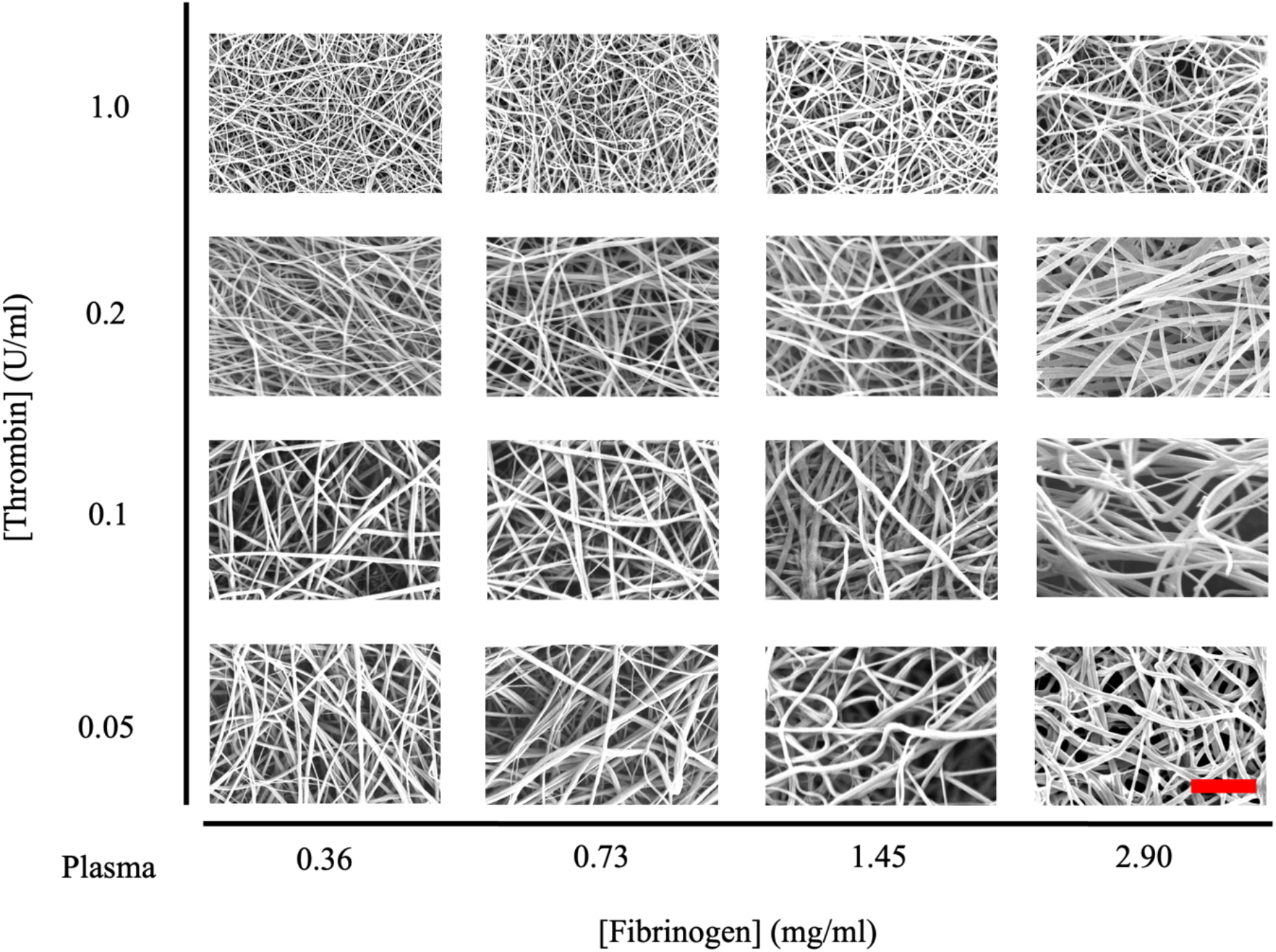
4×4 matrix of representative SEM images showing plasma fibrin network structure across fibrinogen (x-axis) and thrombin (y-axis) concentrations. Each panel corresponds to a specific fibrinogen–thrombin condition. Scale bar: 1 µm.

**Figure 2:**
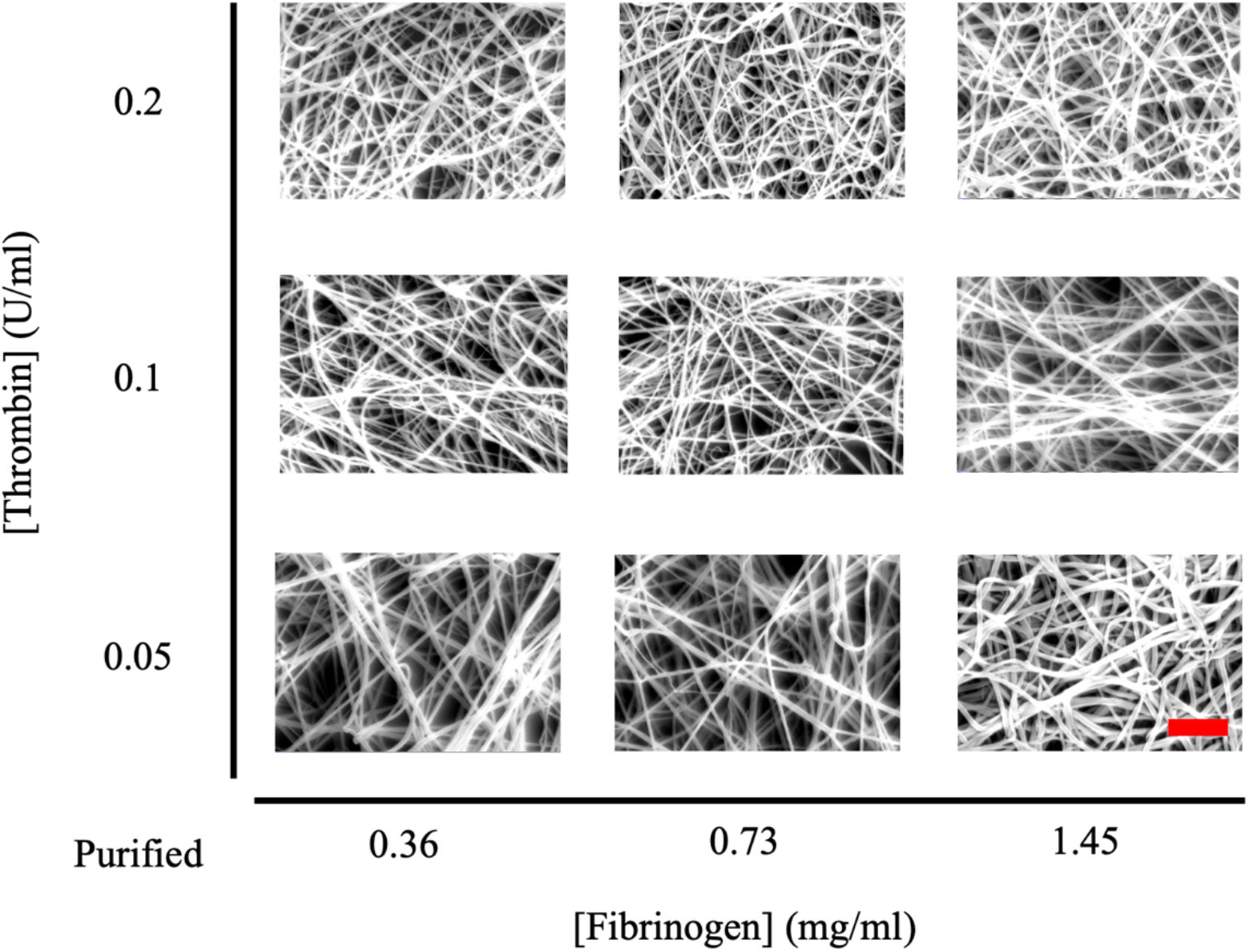
3×3 matrix of representative SEM images showing purified fibrin network structure across fibrinogen (x-axis) and thrombin (y-axis) concentrations. Each panel corresponds to a specific fibrinogen–thrombin condition. Scale bar: 1 µm.

**Figure 3:**
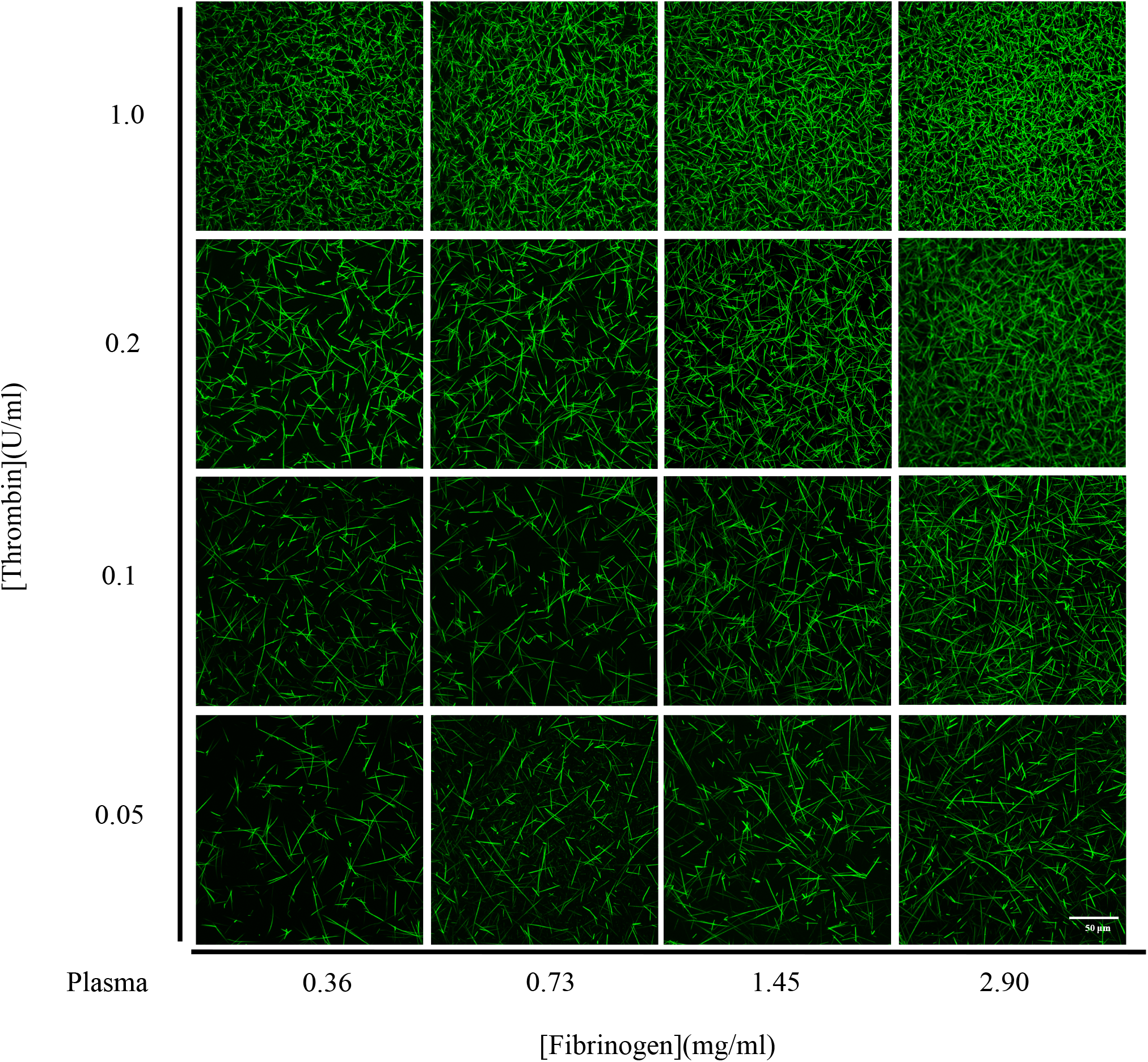
4×4 matrix of representative confocal images showing plasma fibrin network structure across fibrinogen (x-axis) and thrombin (y-axis) concentrations. Each panel corresponds to a specific fibrinogen–thrombin condition. Scale bar: 50 µm.

**Table 1:** Power-law fit parameters and statistical results for plasma fiber diameter, fiber length, fiber density, and bubble diameter using the model *Y* = *k*[*Fgn*]_0_*^α^*[*T*ℎ*r*]_0_*^β^*, with [*Fgn*]_0_ in mg/mL and [*T*ℎ*r*]_0_ in U/mL. The prefactor *k* carries units such that *Y* is in its reported units, i.e., [*k*] = [*Y*] (*mg*/*mL*)^−*α*^(*U*/*mL*)^−*β*^ (fiber diameter in nm, fiber length in µm, fiber density in fibers/1500 µm², and bubble diameter in µm).

| Plasma |  | Value | 95% CI | p-Value | R<sup>2</sup> (log) |
| --- | --- | --- | --- | --- | --- |
| Parameters for Fiber diameter | k | 81 | [80, 85] | <0.001 | 0.96 |
| | $\alpha$ | 0.19 | [0.16, 0.22] | <0.001 | |
| | $\beta$ | -0.23 | [-0.24, -0.21] | <0.001 | |
| Parameters for Fiber length | k | 1155 | [1103, 1211] | <0.001 | 0.96 |
| | $\alpha$ | 0.09 | [0.05, 0.11] | <0.001 | |
| | $\beta$ | -0.24 | [-0.25, -0.21] | <0.001 | |
| Parameters for Fiber density | k | 25 | [23, 25] | <0.001 | 0.88 |
| | $\alpha$ | 0.45 | [0.41, 0.45] | <0.001 | |
| | $\beta$ | 0.30 | [0.29, 0.31] | 0.0015 | |
| Parameters for Bubble diameter | k | 6.6 | [6.5, 6.6] | <0.001 | 0.89 |
| | $\alpha$ | -0.24 | [-0.25, -0.22] | <0.001 | |
| | $\beta$ | -0.14 | [-0.14, -0.13] | <0.001 | |

**Figure 4:**
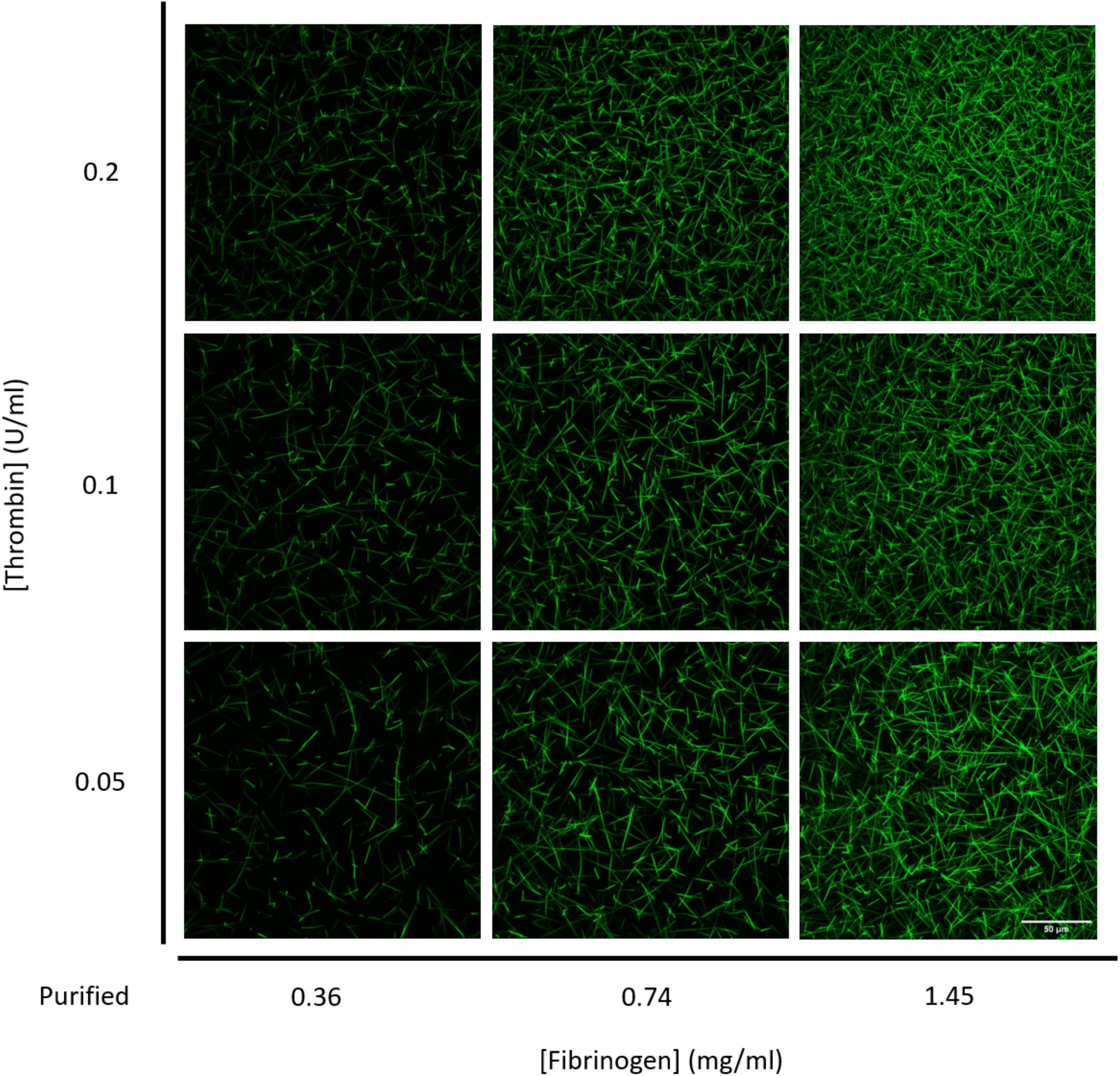
3×3 matrix of representative confocal images showing purified fibrin network structure across fibrinogen (x-axis) and thrombin (y-axis) concentrations. Each panel corresponds to a specific fibrinogen–thrombin condition. Scale bar: 50 µm.

### Power law fits quantify how fibrinogen and thrombin shape fibers and pores

To quantify concentration dependence, each structural metric was fit with a power law *Y* = *k* [*Fgn*]_0_*^α^*[*T*ℎ*r*]_0_*^β^* (Figs. 5 and 6; Tables 1and 2); see the discussion section for a justification for the power law fit. The resulting exponents capture distinct contributions of fibrinogen and thrombin in both plasma and purified systems.

**Figure 5:**
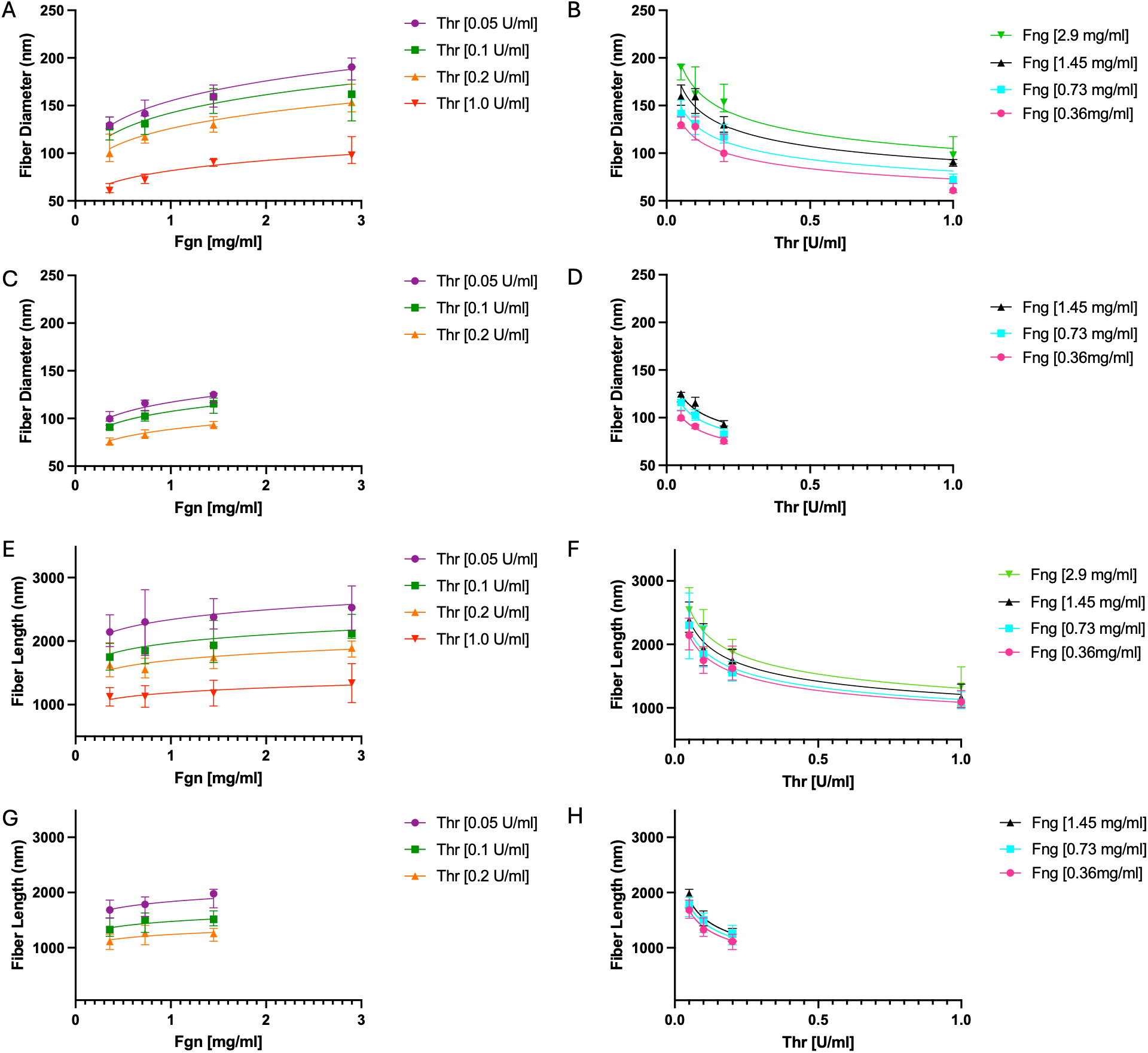
Concentration trends of fiber diameter and fiber length measured from SEM images, with power law fits. For each condition, fiber diameter and fiber length were summarized by the median. Solid lines indicate power law fits. Error bars show 95% confidence intervals across images within each condition. (A,B) Plasma fiber diameter vs fibrinogen and thrombin. (C,D) Purified fiber diameter vs fibrinogen and thrombin. (E,F) Plasma fiber length vs fibrinogen and thrombin. (G,H) Purified fiber length vs fibrinogen and thrombin.

### Fiber diameter (SEM)

Fiber diameter increases with fibrinogen concentration and decreases with thrombin concentration in both plasma and purified fibrinogen clots (Figs. 5; Tables 1 and S1).

### Fiber length (SEM)

Fiber length shows a strong dependence on thrombin (shorter segments at higher thrombin) but weak sensitivity to fibrinogen (Figs. 5; Tables 1 and S2). Consistent with this, the fitted fibrinogen exponent for length is small in both plasma and purified systems (*α* < 0.1; Tables 1 and 2), yielding an almost flat trend versus fibrinogen compared with the thrombin dependence.

### Fiber density and pore size (confocal)

Confocal analysis showed that fiber density increases with both fibrinogen and thrombin, with fibrinogen having the stronger influence (Figs. 6; Tables 2, S3 and S4). Bubble diameter decreased with increasing fibrinogen and thrombin, indicating smaller pore spaces at higher concentrations.

**Figure 6:**
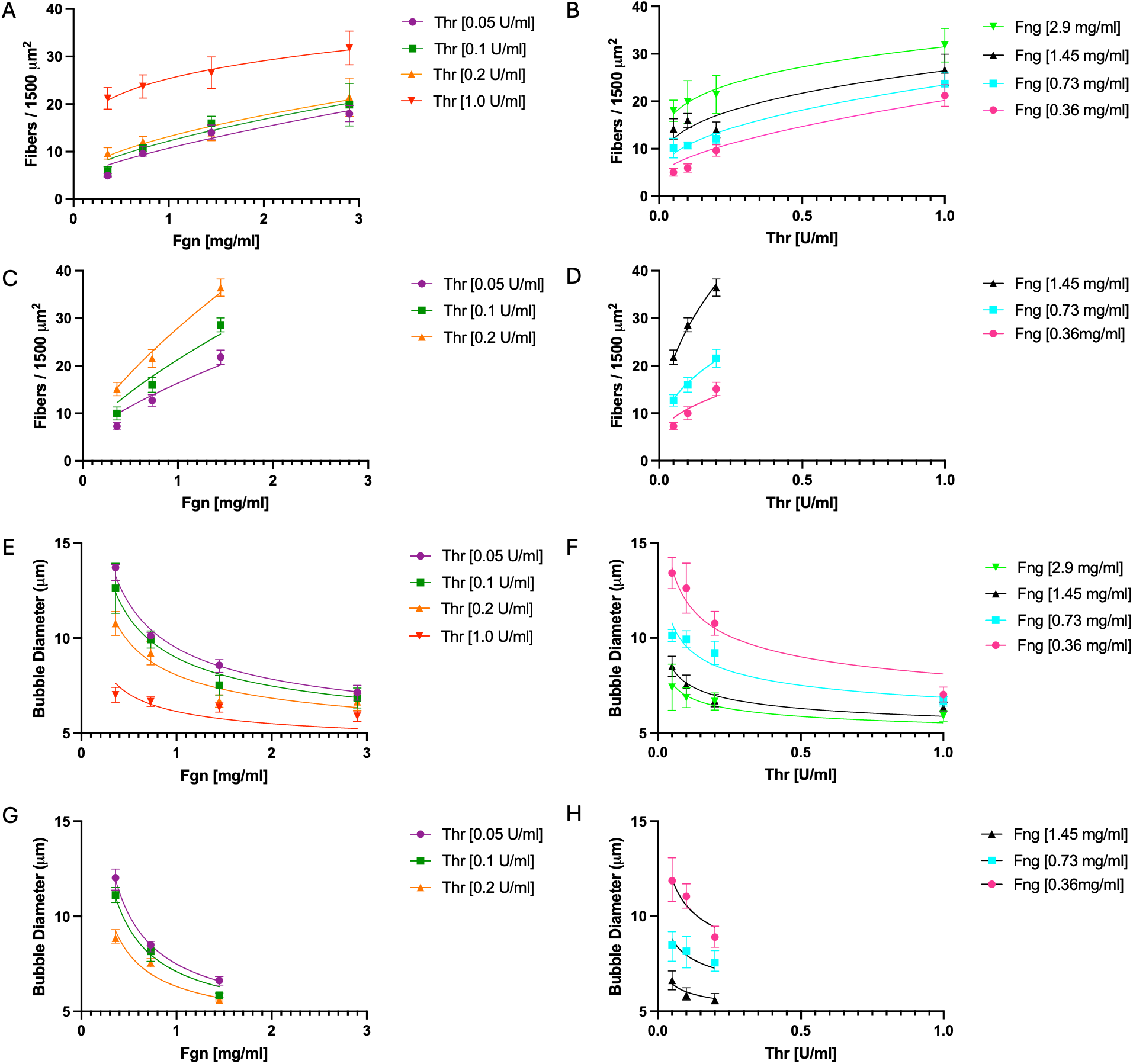
Concentration trends of fiber count and bubble diameter measured from confocal images, with power law fits. For each condition, fiber count was summarized by the mean, whereas bubble diameter was summarized by the median. Solid lines indicate power law fits. Error bars show 95% confidence intervals across images within each condition. (A,B) Plasma fiber count vs fibrinogen and thrombin. (C,D) Purified fiber count vs fibrinogen and thrombin. (E,F) Plasma bubble diameter vs fibrinogen and thrombin. (G,H) Purified bubble diameter vs fibrinogen and thrombin.

## Discussion

### Mechanistic rationale for multiplicative power-law fits

A Michaelis–Menten activation-with-depletion picture provides an intuitive basis for the observed multiplicative power laws (details in Supplement S1). Because the early-time kinetics are complex, we focus on a phenomenological main structure-building window that contributes most strongly to the observed fibrin network structure. During this window we make a simple balance approximation: the supply of incorporation-competent fibrin species and their incorporation into growing structures are of similar magnitude, so the size of this “activated” intermediate pool stays roughly constant (production ≈ consumption). Depletion then drives the effective fibrinogen level in this window into the low-substrate limit ([*Fgn*] ≪ *K_m_*), making the Michaelis–Menten term approximately linear in [*Fgn*]. Together, this yields power-law scaling of effective growth interfaces with [Fgn]_0_ and [Thr]_0_, and a geometric mass-allocation argument translates that scaling into the observed power-law dependence of single-fiber metrics.

### Conserved fibrinogen and thrombin concentration sensitivities, and environment-dependent scaling (plasma vs purified system)

Across plasma and purified systems, the fitted power-law exponents for single-fiber diameter and length are similar, indicating that the core mechanisms governing single-fiber growth (elongation and lateral aggregation) are regulated by [*Fgn*]_0_ and [*T*ℎ*r*]_0_ in a largely conserved manner across both systems. In contrast, the prefactor *k* differs between the two systems (by ∼ 1.28 × for fiber diameter, ∼ 1.49 × for fiber length, and ∼ 1.30 × for bubble diameter, with an opposite-direction ∼ 2.26 ×difference for fiber density). Because effective Ca^2+^ and NaCl concentrations were similar, a possible simple explanation for the change in *k* is reduced effective thrombin activity in plasma due to thrombin consumption and inhibition (e.g., antithrombin, with contributions from *α*_2_-macroglobulin and heparin cofactor II). Consistent with reduced effective thrombin activity in plasma [23], a turbidity study reported markedly faster polymerization kinetics in purified fibrinogen compared to plasma under comparable conditions [12]. If we model this as a simple rescaling *T*_eff_ = *sT*_0_, then *k*_plasma_ = *k*_pur_*s^β^*, giving

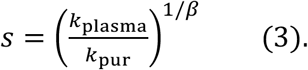

Applying this estimate to SEM-derived single-fiber metrics (diameter and length) yields *s* ≈ 0.19–0.35, i.e., [*T*ℎ*r*]_eff_ is reduced by ∼ 65–81% in plasma.

### Kinetic control (thrombin) vs space-filling control (fibrinogen)

Our scaling analysis elucidates the different roles of thrombin and fibrinogen: thrombin primarily sets the gelation dynamics and thus the effective growth for individual fibers, whereas fibrinogen primarily sets how the network fills space. Mechanistically, increasing thrombin accelerates fibrinogen to fibrin conversion and early assembly, effectively increasing the formation rate of assembly-competent species and the density of nucleation sites (“seeds”). A higher density of nucleation sites means protofibrils, fibrin aggregates and early fibers encounter and connect with each other sooner, thereby advancing the percolation/gel point and shortening the time available for continued elongation of any single fiber. Consistent with this kinetic picture, fiber length is overwhelmingly thrombin-set—doubling thrombin *shortens* length by 15% in plasma (2^-0.24^ = 0.85) and 18% in purified fibrinogen (2^-0.29^ = 0.82), while doubling fibrinogen *increases* length by only 5–7% (2^0.07^–2^0.09^ = 1.05–1.07)—arguing that elongation is limited by thrombin-controlled fiber progression rather than fibrinogen monomer-limited. In the same framework, a larger number of nucleation sites yields a higher fiber density, and the faster kinetic partitioning of a finite material supply among more growing units biases fibers toward being slightly thinner. Indeed, fiber density increases with thrombin (β = 0.30 in plasma; β = 0.43 in purified fibrinogen), while fiber diameter decreases with thrombin (β = - 0.23 and - 0.21, respectively).

In contrast, fibrinogen strongly influences the space-filling architecture: doubling fibrinogen increases fiber density by 37% in plasma (2^0.45^ = 1.37) and 66% in purified fibrinogen (2^0.73^ = 1.66), while reducing bubble diameter by 15% in plasma (2^-0.24^ = 0.85) and 25% in purified fibrinogen (2^-0.41^ = 0.75), respectively. Thrombin’s effect on bubble diameter is comparatively small (9% per doubling; 2^-0.14^ = 0.91 in both systems). Notably, the fibrinogen exponent for bubble diameter is approximately half (with opposite sign) of the fibrinogen exponent for fiber density in both systems (−0.24/0.45 = −0.53; −0.41/0.73 = −0.56), implying a near-geometric packing relation where bubble diameter 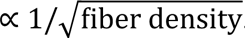. The weaker density–fibrinogen scaling in plasma compared to purified fibrinogen (α = 0.45 vs 0.73) could reflect that at high fiber densities (high fibrinogen), fiber density may be underestimated because of overlapping fibers. A similar effect may contribute to the weaker bubble–fibrinogen scaling in plasma compared to purified fibrinogen (α = −0.24 vs −0.41). In support of this notion is that when fitting identical 3x3 matrices of plasma sample and purified fibrinogen samples, the exponents are the same for densities and bubble diameter. Finally, fiber diameter depends oppositely on fibrinogen and thrombin concentrations; increasing fibrinogen thickens fibers (+11–14% per doubling) while increasing thrombin thins fibers (−14–15% per doubling); so, these effects can largely cancel, yielding a weak net diameter dependence even as fiber density and bubble size shift dramatically.

### Space filling (volume fraction) is superlinear with fibrinogen and contracts with thrombin

The fiber volume fraction is defined as the ratio of the volume occupied by all fibrin fibers to the total observation volume,

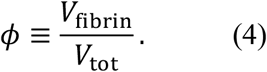

Approximating each fiber as a cylinder of diameter *d* and effective length *l*, the volume per fiber is (*π*/4)*d*^2^*l*. If *N* is the total number of fibers in the observation volume, then

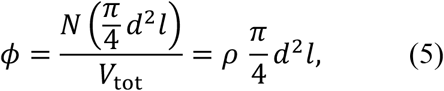

where *ρ* ≡ *N*/*V*_tot_ is the 3D fiber number density.

In our experiments we directly measure the projected (2D) fiber density

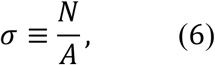

where *A* is the projected imaging area. For an isotropic, space-filling network, dimensional consistency implies σ = *ρ*^2/3^, and hence

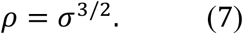

Combining Eqs. (5)–(7) gives an expression for the volume fraction in terms of measured quantities,

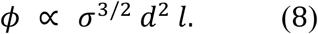

Equation (8) contains only quantities we measure (*σ*, *d*, *l*), so it directly enables an empirical test of how the fiber volume fraction scales with fibrinogen and thrombin.

Under the simplest “mass supply” expectation — homogeneous internal fiber mass density (note, this fiber mass density = (fiber mass)/(fiber volume), is different from fiber number density = (# of fibers)/(volume)) and conversion set primarily by fibrinogen availability — one would anticipate *φ* to increase linearly with [*Fgn*]_0_ (more fibrinogen ⇒ more fibrin volume) and to be independent of [*T*ℎ*r*]_0_ (thrombin reshapes morphology but does not itself contribute fiber volume). Writing the measured power-law scalings, 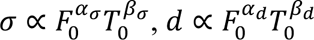, and 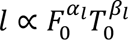, Eq. (8) implies

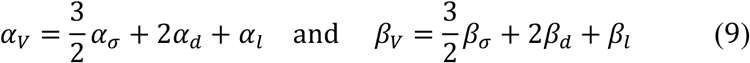

where 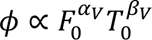

Using the fitted exponents in Table 1 yields *α_V_* = 1.147 and *β_V_* = −0.25 in plasma, and *α_V_* = 1.466 and *β_V_* = −0.07 in purified fibrinogen. If increasing fibrinogen simply increased fiber volume at fixed fiber mass density, one would expect *α_V_* ≃ 1. Instead, *α_V_* > 1, consistent with fibrinogen driving a shift toward thicker fibers accompanied by reduced internal fiber mass density (thicker fibers have lower mass density), as observed previously [24]. In contrast, *β_V_* is close to zero in purified fibrinogen, supporting the idea that thrombin primarily redistributes morphology/partitioning of single fibers rather than increasing or decreasing the total fiber volume fraction. The negative *β_V_* in plasma indicates that a shift toward thinner fibers is accompanied by increased fiber mass density (thinner fibers have higher mass density). Alternatively, if fiber density has been somewhat underestimated in plasma (for the highest fibrinogen concentration), a correction for this underestimation would yield *β_V_* of about zero also in plasma.

### Why is fiber segment length mainly dependent on thrombin? Agreement between fiber segment length and space per thrombin molecule suggests a one-fiber-per-one-thrombin-molecule picture

To relate the measured 3D branch-to-branch segment length, *L*_seg_, to a spatial control parameter, we converted thrombin concentration to a thrombin molecular number density, *ρ*_Thr_. For the used lot, 1 NIH U/mL corresponds to 9 nM thrombin, giving *ρ*_Thr_ ≈ 5.42 *μ*m^−3^ at 1 U/mL, meaning there are 5.42 thrombin molecules per 1 *μ*m^−3^ (and a *ρ*_Thr_ of 1.08, 0.542, and 0.271 *μ*m^−3^ for 0.2, 0.1, and 0.05 U/mL thrombin, respectively). From *ρ*_Thr_, we compute the mean volume per thrombin molecule, *V*_part_ = 1/*ρ*_Thr_. Representing *V*_part_ as a cube provides a simple geometric spacing scale: the cube side is 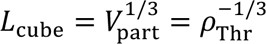, and the cube diagonal is *L*_diag_ = 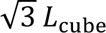. If segments are approximately straight over these length scales, then *L*_cube_ and *L*_diag_ provide lower and upper bounds on the center-to-center spacing expected from the thrombin number density, so [*L*_cube_, *L*_diag_] serves as a thrombin-implied spacing range.

We compared this spacing range (the space occupied by one thrombin molecule) to the measured *L*_seg_ in both plasma and purified fibrinogen clots. Because fibrinogen has only a minor effect on *L*_seg_, we pooled *L*_seg_ values across fibrinogen conditions at a fixed thrombin level and compared the pooled *L*_seg_ range to [*L*_cube_, *L*_diag_] for that thrombin level (Table 3). Strikingly, across all thrombin concentrations, the pooled *L*_seg_ values lie in the same micrometer range as [*L*_cube_, *L*_diag_], and both contract systematically as thrombin increases. This agreement supports a direct interpretation: at a given thrombin concentration, the average volume “assigned” to one thrombin molecule contains on the order of one fiber segment, so the characteristic segment length is set primarily by the spacing between individual thrombin molecules.

**Table 2:** Power-law fit parameters and statistical results for purified fiber diameter, fiber length, fiber density, and bubble diameter using the model *Y* = *k*[*Fgn*]_0_*^α^*[*T*ℎ*r*]_0_*^β^*, with [*Fgn*]_0_ in mg/mL and [*T*ℎ*r*]_0_ in U/mL. The prefactor *k* carries units such that *Y* is in its reported units, i.e., [*k*] = [*Y*] (*mg*/*mL*)^−*α*^(*U*/*mL*)^−*β*^ (fiber diameter in nm, fiber length in µm, fiber density in fibers/1500 µm², and bubble diameter in µm).

| Purified Fibrinogen |  | Value | 95% CI | p-Value | R<sup>2</sup> (log) |
| --- | --- | --- | --- | --- | --- |
| Parameters for Fiber diameter | k | 64 | [60., 66] | <0.001 | 0.96 |
| | $\alpha$ | 0.15 | [0.14, 0.18] | <0.001 | |
| | $\beta$ | -0.21 | [-0.24, -0.20] | <0.001 | |
| Parameters for Fiber length | k | 775 | [688, 867] | <0.001 | 0.84 |
| | $\alpha$ | 0.07 | [0.04, 0.14] | 0.0057 | |
| | $\beta$ | -0.29 | [-0.34, -0.24] | 0.0084 | |
| Parameters for Fiber density | k | 57 | [54, 58] | <0.001 | 0.99 |
| | $\alpha$ | 0.73 | [0.70, 0.74] | <0.001 | |
| | $\beta$ | 0.43 | [0.40, 0.44] | <0.001 | |
| Parameters for Bubble diameter | k | 5.1 | [4.9, 5.2] | <0.001 | 0.97 |
| | $\alpha$ | -0.41 | [-0.42, -0.39] | <0.001 | |
| | $\beta$ | -0.14 | [-0.15, 0.13] | 0.0029 | |

**Table 3:** Thrombin number density and implied spacing scale compared with measured 3D fibrin segment length. (particles·*μ*m^−3^) denotes thrombin molecular number density derived from thrombin activity (NIH U · mL^−1^; parentheses; lot activity 3044 NIH U · mg^−1^). The spacing range [*L*_cube_, *L*_diag_] is defined by 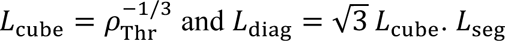 and *L*_diag_ = √3 *L*_cube_. *L*_seg_ ranges (plasma and purified fibrinogen) are pooled across fibrinogen conditions at fixed thrombin.

| Thrombin Density<br>(particles/$\mu\text{m}^3$) | Thrombin Space Range<br>$L_{\text{cube}}$ ($\mu\text{m}$), $L_{\text{diag}}$ ($\mu\text{m}$) | Plasma<br>$L_{\text{seg}}$ range ($\mu\text{m}$) | Purified Fibrinogen<br>$L_{\text{seg}}$ range ($\mu\text{m}$) |
| --- | --- | --- | --- |
| 5.42 (1 U/mL) | [0.57, 0.99] | [1.09, 1.30] |  |
| 1.08 (0.2 U/mL) | [0.97, 1.69] | [1.55, 1.89] | [1.11, 1.26] |
| 0.542 (0.1 U/mL) | [1.23, 2.12] | [1.75, 2.12] | [1.33, 1.52] |
| 0.271 (0.05 U/mL) | [1.55, 2.68] | [2.14, 2.53] | [1.69, 1.98] |

An order-of-magnitude kinetics check supports the plausibility of this thrombin-spacing picture. Using an approximate fibrinogen molecular weight of 340 kDa, our fibrinogen range (0.36–2.9 mg/mL) corresponds to ∼1 – 9 µM, or ∼6×10² – 5×10³ molecules·µm⁻³. Combined with the thrombin number densities above, the mean volume per thrombin molecule contains on the order of 10³ – 10⁴ fibrinogen molecules at 0.05–0.1 U/mL. Reported catalytic efficiencies for thrombin-catalyzed fibrinopeptide A release are ∼10⁷ M⁻¹ s⁻¹ [25], implying per-enzyme turnover rates of tens of s⁻¹ over our fibrinogen range. These values indicate that a single thrombin molecule (or a small local cluster acting as an effective center) could activate the fibrinogen molecules within its assigned volume on a timescale of minutes, consistent with the assembly window of our experiments.

One plausible mechanism is a “one-nucleation center-per-thrombin” picture in which each thrombin molecule (or a small local cluster acting as one effective center) locally generates a burst of assembly-competent fibrin that accumulates within its assigned volume and builds a short segment. Gelation then occurs as these locally formed segments elongate, thicken, fill space, and encounter neighboring segments; once segments meet, they connect and the network becomes mechanically locked into a percolated scaffold. In this view, the typical segment length is limited by the thrombin-set spacing between neighboring initiation regions, giving a simple geometric scaling 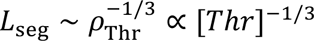. By contrast, fibrinogen primarily modulates how much fibrin mass is available within each initiation volume — thereby affecting thickness, mass-per-length, and space-filling — without strongly shifting the characteristic segment-length scale set by thrombin spacing.

### Advances and limitations

#### Scope and robustness of the scaling analysis

Although related questions have been explored previously using similar experimental modalities, the present study provides a substantially more comprehensive and internally consistent characterization of how fibrinogen and thrombin jointly shape fibrin architecture across both plasma and purified systems. Specifically, we systematically mapped a 4×4 matrix in human plasma and a 3×3 matrix in purified fibrinogen, enabling direct comparisons across a wide range of [*Fgn*]–[*T*ℎ*r*] combinations. Prior work typically examined a much smaller subset of conditions — for example, varying thrombin at fixed fibrinogen or vice versa — often yielding weak or purely phenomenological linear fits with limited explanatory power [12]. Importantly, our cross-system comparisons (plasma vs. purified fibrinogen) were performed at matched thrombin concentrations in plasma and purified fibrinogen, avoiding confounds introduced by using different initiators (e.g., thrombin in one system versus tissue factor in the other), which can obscure mechanistic interpretation [12].

To assess the uncertainty and robustness of the fitted power-law relationships, we examined the fitted parameters k, α, and β using 2000-iteration bootstrap resampling to calculate 95% confidence intervals, which are reported in Tables 1 and 2. These intervals were closely centered around the fitted parameter estimates, supporting the stability of the fitted k, α, and β values. We further performed leave-one-condition-out refitting, in which each fibrinogen–thrombin concentration pair was omitted in turn and the model was refit. As summarized in Supplementary Tables S5 and S6, the fitted α and β values changed only modestly across these refits, supporting the conclusion that the observed trends were not dominated by any single concentration condition.

In this power-law formulation, the fitted exponents may be interpreted as logarithmic sensitivity, or elasticity coefficients: α = ∂lnY/∂ln[Fgn] and β = ∂lnY/∂ln[Thr]. They quantify the fractional change in each structural metric associated with a fractional change in fibrinogen or thrombin concentration within the fitted experimental range.

#### SEM-based quantification of single-fiber dimensions

A key technical advance is that single-fiber dimensions were quantified using a standardized SEM preparation and imaging protocol, providing high-resolution, quantitatively robust measurements of fiber diameter and branch-to-branch segment length. Optical approaches can reveal qualitative trends in diameter but are intrinsically limited by diffraction and labeling/thresholding artifacts, especially for submicron fibers, making diameter estimates approximate [14]. By contrast, SEM under a standardized protocol resolves fiber boundaries with sufficient precision to support reliable comparisons across conditions.

Segment length has been particularly challenging to quantify with existing microscopy-based workflows [15]. Here we leveraged SEM at moderate magnification and analyzed only the central portion of each image to minimize edge truncation, while maintaining a short working distance to ensure that measured segments lie within a well-defined focal plane. This combination reduces common biases in segment-length measurements, especially those arising from fibers leaving the field of view or from depth ambiguity and enables precise quantification of *L*_seg_.

These considerations may also help explain discrepancies with prior reports based on 2D confocal single-plane measurements that concluded segment length decreases with increasing fibrinogen and thrombin [12]. In that study, increasing fibrinogen was reported to shorten apparent segment length, whereas we observe the opposite trend (slight lengthening). At high network density, limited resolution and ambiguous identification of true branch points versus projected crossings or fiber pile-up can systematically bias 2D optical measurements toward shorter inferred segments. In contrast, SEM can provide high-resolution surface-topographic information that allows many intersections and branch points in the topmost resolved fiber layer to be distinguished based on local morphology. By excluding ambiguous crossings and truncated or out-of-focus segments, our SEM-based analysis provides a more conservative and reliable assessment of how fibrinogen and thrombin modulate apparent branch-to-branch segment length, *L*_seg_ , across plasma and purified fibrinogen systems.

#### Implications for fibrin mechanics

These structural trends are also relevant to the viscoelastic behavior of fibrin gels. Prior studies have shown that bulk clot elastic or storage modulus generally increases with fibrinogen concentration, although the reported dependence varies across systems and experimental conditions, ranging from near-linear behavior over some tested concentration ranges to power-law scaling [9, 24, 26]. At the single-fiber level, fibrin fiber modulus decreases with increasing fiber diameter, approximately as a power law [24]. Together, these findings indicate that fibrin mechanics emerge from coupled contributions of single-fiber properties and network architecture, rather than from any single structural parameter alone.

In this context, our scaling results suggest that fibrinogen and thrombin influence clot mechanics through distinct but overlapping structural pathways. Fibrinogen has its strongest effects on space-filling features such as fiber density and pore/bubble size, while also contributing to fiber thickening, whereas thrombin more strongly regulates branch-to-branch segment length and network connectivity, while also tending to reduce fiber diameter. However, because we did not directly measure rheological properties in the present study, we refrain from making quantitative predictions for bulk modulus and instead view the present scaling relations as a structural framework for future structure–mechanics studies.

#### Interpretation and limitations of confocal network measurements

Network-level metrics derived from confocal microscopy, including fiber density and pore/bubble readouts, are subject to important limitations. Because the analysis relies on resolvable features in confocal images and on projections/binarization, these quantities should be interpreted as resolution-limited trends rather than absolute 3D ground-truth values, particularly in the densest networks where overlap can lead to undercounting fiber density or merged pores. Consistent with this limitation, we reduced the purified fibrinogen clots condition matrix from 4×4 to 3×3, as clots formed at the highest fibrinogen and thrombin concentrations were too dense to quantify reliably without introducing bias that could confound cross-condition comparisons.

Fluorescent labeling of fibrinogen may also impose constraints on confocal-based structural measurements. In the present study, Alexa Fluor 488-labeled fibrinogen was used at 6.5% of the total fibrinogen concentration. This fraction was selected empirically to provide sufficient fluorescence signal across the 212 μm × 212 μm × 15 μm imaging volume while avoiding obvious label-associated aggregation or star-like fluorescent features observed at higher labeling levels. Prior work has shown that fluorescently labeled fibrinogen can affect fibrin formation dynamics and fibrin structure, as measured by turbidity [24]. To further evaluate this effect under our experimental conditions, turbidity measurements were performed in both plasma and purified fibrinogen systems at fibrinogen concentrations of 0.36, 0.72, 1.45, and 2.4 mg/mL using 0, 1.5, 3, 6, and 12% Alexa Fluor 488-labeled fibrinogen (Figures S1 and S2). Minimal differences in fibrin formation dynamics and final turbidity were observed up to 6% labeling, whereas more noticeable deviations appeared at 12% labeling. This is consistent with the appearance of star-like fluorescent aggregates observed in confocal imaging at labeling fractions above approximately 6.5%, suggesting that excessive labeling may promote local aggregation or alter fibrin assembly. Therefore, although subtle effects of labeling cannot be completely excluded, the 6.5% labeling fraction used in this study was chosen as a practical compromise that maintained sufficient fluorescence intensity for reproducible quantification of fiber density and pore/bubble size while minimizing detectable perturbations to fibrin network formation and structure.

More broadly, the fitted scaling relations are supported within the experimentally resolved range of fibrinogen (0.36–2.9 mg/mL) and thrombin (0.05–1 U/mL), where robust clot formation and quantitative imaging were possible. They have not been validated quantitatively outside this range. Below this fibrinogen range, stable clot formation was difficult, and network architecture was therefore not well defined. Below this thrombin range, particularly at low fibrinogen concentrations, clot formation became extremely slow, with mature networks requiring hours or longer to develop. At higher fibrinogen concentrations, increasing network density made quantitative structural characterization challenging, particularly for confocal-based metrics. At higher thrombin concentrations, clotting was too rapid to allow reproducible pipetting and sample handling without perturbing the nascent network.

Confocal imaging was performed at room temperature. Temperature can influence fibrin polymerization kinetics, with physiological temperature accelerating early fibrinopeptide release and clot formation relative to room temperature. However, prior work suggests that temperature primarily affects the rate of polymerization rather than fundamentally altering the pre-gel polymerization pathway [27], and a recent comparison of clots formed at room temperature versus 37 °C found that temperature-dependent structural differences were smaller than those caused by changes in fibrinogen or thrombin concentration [12]. Thus, although small quantitative shifts in fiber density or pore/bubble measurements at 37 °C cannot be excluded, we do not expect physiological temperature to qualitatively alter the concentration-dependent trends reported here.

Despite these limitations, confocal imaging provides an important advantage for network-level readouts such as fiber density and pore size: it quantifies the clot in a hydrated, near-native state, minimizing dehydration- and shrinkage-related distortions that can strongly affect apparent network compaction under electron-microscopy sample preparation.

#### Future directions and experimental considerations

Some limitations of confocal microscopy may be addressed in future studies using complementary high-resolution fluorescence methods. Super-resolution approaches such as STED, PALM, and STORM could improve separation of closely spaced fibrin fibers in selected dense regions and reduce errors caused by unresolved fiber overlap. However, for large-volume, quantitative mapping of thick, hydrated fibrin networks, these approaches would need to be balanced against practical considerations such as imaging depth, optical scattering or aberration, acquisition time, photobleaching, field of view, and fluorophore/labeling requirements. Thus, super-resolution microscopy may provide valuable validation of selected conditions or local network regions, while confocal microscopy remains useful for hydrated, large-volume network-level measurements such as fiber density and pore/bubble size.

Ca²⁺ concentration is another variable that can influence fibrin polymerization and network architecture. In plasma clots, CaCl₂ was required to overcome citrate anticoagulation and initiate clot formation. In purified fibrinogen clots, where citrate was absent, CaCl₂ was added under the matched fixed-calcium condition used for comparison with the recalcified plasma experiments, while also helping low-fibrinogen or low-thrombin samples clot within a practical experimental time window. Because CaCl₂ was held constant across each fibrinogen–thrombin matrix, the fitted α and β values should be interpreted as concentration-dependent trends under this fixed calcium condition.

## Conclusion

Fibrin architecture emerges from coupled enzyme activation, substrate depletion, formation of growth interfaces, and geometric mass allocation, yet predicting how fibrinogen and thrombin jointly shape microstructure remains challenging. By mapping fibrin structure across fibrinogen– thrombin concentration matrices in human plasma and purified fibrinogen, we found that a broad set of structural readouts is well described by compact multiplicative power laws *Y* = *k*[*Fgn*]_0_*^α^*[*T*ℎ*r*]_0_*^β^*. We present a simplified, coarse-grained kinetic rationale (Michaelis–Menten activation with depletion and a quasi-steady reduction during the main assembly window) to motivate why separable, power-law-like sensitivities can emerge over a finite experimental range.

The exponent patterns expose a division of labor between thrombin and fibrinogen. Thrombin acts primarily as a single-fiber kinetic controller: increasing thrombin accelerates early assembly and progression to connectivity, compressing the effective growth window and shortening the characteristic branch-to-branch segment length, consistent with a spatial “territory” interpretation in which segment length contracts on a thrombin-set micrometer scale. Fibrinogen acts primarily as a space-filling controller, strongly increasing fiber density and reducing pore/bubble size while thickening fibers, with only weak influence on segment length. At the network scale, the fitted exponents further imply an emergent, isotropic packing rule close to 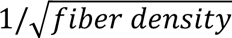, providing a simple geometric bridge between connectivity and transport-relevant void structure.

Finally, plasma and purified systems exhibited similar sensitivities (exponents), suggesting a shared underlying fiber assembly mechanism, while the dominant system difference appeared as a shift in the prefactor *k*, consistent with environment-dependent effective thrombin activity and regulation in plasma. Together, these results provide a predictive scaling atlas and a mechanistically interpretable separation of roles — thrombin setting single-fiber kinetic/length scales and fibrinogen tuning space-filling architecture — useful for theory development and for interpreting how clinically relevant shifts in fibrinogen or thrombin may reshape clot microstructure. Because fibrin microstructure helps determine clot-level function, these concentration-dependent structural changes may have implications beyond morphology alone. Changes in fiber density, pore size, fiber diameter, and network connectivity can influence clot permeability, mechanical stability, and susceptibility to fibrinolysis. For example, denser networks with smaller pores may restrict transport of fibrinolytic enzymes and alter lysis behavior, whereas changes in fiber thickness and connectivity may affect clot stability and fragmentation. Thus, the scaling relationships reported here provide a structural framework for future studies linking changes in coagulation-factor levels to fibrinolysis, hemostatic balance, and thrombotic or cardiovascular risk. This framework can also be extended to examine how additional plasma components, cellular elements, crosslinking, platelets, inflammatory factors, or therapeutic agents further modify fibrin architecture.

## Declaration of AI and AI-assisted technologies in the writing process

During the preparation of this work the author(s) used ChatGPT (OpenAI) to improve English grammar and clarity. After using this tool, the author(s) reviewed and edited the content as needed and take full responsibility for the content of the published article.

## Author contributions

Conceptualization: CC, ZZ, AN, DN, KB, BEB, NEH, SRB, MG

Data curation: CC, ZZ, AN, DN, KB, BEB, NEH, GM, SRB, MG

Formal analysis: CC, ZZ, AN, DN, KB, SRB, MG

Funding acquisition: NEH, BEB, MG

Investigation: CC, ZZ, AN, DN, KB, BEB, NEH, GM, SRB, MG

Methodology: CC, ZZ, AN, DN, KB, BEB, NEH, GM, SRB, MG

Project administration: NEH, BEB, MG

Resources: NEH, BEB, MG

Software: CC, ZZ, AN, DN, KB

Supervision: MG

Validation: CC, ZZ, AN, KB, BEB, NEH, SRB, MG

Visualization: CC, ZZ, AN, SRB

Roles/Writing - original draft: CC, ZZ, SRB, MG

and Writing - review & editing: CC, ZZ, AN, KB, BEB, NEH, SRB, MG

## Acknowledgements

This work was supported by National Institutes of Health grants 2R15HL148842-02. The content is solely the responsibility of the authors and does not necessarily represent the official views of the National Institutes of Health. We thank Xiao Ma and Dundappa Mumbaraddi from the Department of Chemistry at Wake Forest University for their assistance with SEM imaging.

## References

[1] J.W. Weisel, R.I. Litvinov, Mechanisms of fibrin polymerization and clinical implications, Blood 121(10) (2013) 1712–1719.

[2] A. Undas, R.A.S. Ariëns, Fibrin Clot Structure and Function A Role in the Pathophysiology of Arterial and Venous Thromboembolic Diseases, Arterioscl Throm Vas 31(12) (2011) E88–E99.

[3] J. Danesh, S. Lewington, S.G. Thompson, G.D.O. Lowe, R. Collins, F.S. Collaboration, Plasma fibrinogen level and the risk of major cardiovascular diseases and nonvascular mortality - An individual participant meta-analysis, Jama-J Am Med Assoc 294(14) (2005) 1799–1809.

[4] J. Klovaite, B.G. Nordestgaard, A. Tybjærg-Hansen, M. Benn, Elevated Fibrinogen Levels Are Associated with Risk of Pulmonary Embolism, but Not with Deep Venous Thrombosis, Am J Resp Crit Care 187(3) (2013) 286–293.

[5] B. Charbit, L. Mandelbrot, E. Samain, G. Baron, B. Haddaoui, H. Keita, O. Sibony, D. Mahieu-Caputo, M.F. Hurtaud-Roux, M.G. Huisse, M.H. Denninger, D. De Prost, P.S. Grp, The decrease of fibrinogen is an early predictor of the severity of postpartum hemorrhage, J Thromb Haemost 5(2) (2007) 266–273.

[6] R. Sidonio, M. Hoffman, G. Kenet, Y. Dargaud, Thrombin generation and implications for hemophilia therapies: A narrative review, Res Pract Thromb Hae 7(1) (2023) e100018, 1-11.

[7] C.J. Reddel, C.W. Tan, V.M. Chen, Thrombin Generation and Cancer: Contributors and Consequences, Cancers 11(1) (2019) 100, 1-20.

[8] R. Gehlen, A. Vandevelde, B. de Laat, K.M.J. Devreese, Application of the thrombin generation assay in patients with antiphospholipid syndrome: A systematic review of the literature, Front Cardiovasc Med 10 (2023) 1075121, 1-17.

[9] E.A. Ryan, L.F. Mockros, J.W. Weisel, L. Lorand, Structural origins of fibrin clot rheology, Biophysical Journal 77(5) (1999) 2813–2826.

[10] F. Ferri, M. Greco, G. Arcòvito, M. De Spirito, M. Rocco, Structure of fibrin gels studied by elastic light scattering techniques: Dependence of fractal dimension, gel crossover length, fiber diameter, and fiber density on monomer concentration, Phys Rev E 66(1) (2002) 011913.

[11] H.A. Belcher, M. Guthold, N.E. Hudson, What is the diameter of a fibrin fiber?, Res Pract Thromb Hae 7(5) (2023) e100285, 1-11.

[12] R.A. Risman, H.A. Belcher, R.K. Ramanujam, J.W. Weisel, N.E. Hudson, V. Tutwiler, Comprehensive Analysis of the Role of Fibrinogen and Thrombin in Clot Formation and Structure for Plasma and Purified Fibrinogen, Biomolecules 14(2) (2024) 230, 1-18.

[13] J.W. Weisel, Structure of fibrin: impact on clot stability, J Thromb Haemost 5 (2007) 116–124.

[14] K. Pietsch, L. Storm-Johannsen, A. Schmidt-Thomée, T. Pompe, Correlation between Fibrin Fibrillation Kinetics and the Resulting Fibrin Network Microstructure, Adv Healthc Mater 12(8) (2023) 2202231, 1-11.

[15] J.W. Weisel, C. Nagaswami, Computer Modeling of Fibrin Polymerization Kinetics Correlated with Electron-Microscope and Turbidity Observations - Clot Structure and Assembly Are Kinetically Controlled, Biophysical Journal 63(1) (1992) 111–128.

[16] O.V. Kim, R.I. Litvinov, J.W. Weisel, M.S. Alber, Structural basis for the nonlinear mechanics of fibrin networks under compression, Biomaterials 35(25) (2014) 6739–6749.

[17] C. Cai, Z. De Lange-Loots, S.R. Baker, R.I. Litvinov, A.C. Swanepoel, C. Nagaswami, J.W. Weisel, R. Marchi, A. Casini, M. Neerman-Arbez, C. Duval, R.A.S. Ariens, R.A. Risman, V. Tutwiler, H.A. Belcher, A. Elangovan, N.E. Hudson, A. Undas, M. Zabczyk, M. Guthold, M. Pieters, Standardization of scanning electron microscopy analysis of fibrin fiber diameter measurement: communication from the ISTH SSC subcommittee on FXIII and fibrinogen, J Thromb Haemost 24(1) (2026) 318–328.

[18] M. Molteni, D. Magatti, B. Cardinali, M. Rocco, F. Ferri, Fast Two-Dimensional Bubble Analysis of Biopolymer Filamentous Networks Pore Size from Confocal Microscopy Thin Data Stacks, Biophysical Journal 104(5) (2013) 1160–1169.

[19] W. Mickel, S. Münster, L.M. Jawerth, D.A. Vader, D.A. Weitz, A.P. Sheppard, K. Mecke, B. Fabry, G.E. Schröder-Turk, Robust Pore Size Analysis of Filamentous Networks from Three-Dimensional Confocal Microscopy, Biophysical Journal 95(12) (2008) 6072–6080.

[20] C.L. Chiu, V. Hecht, H. Duong, B. Wu, B. Tawil, Permeability of Three-Dimensional Fibrin Constructs Corresponds to Fibrinogen and Thrombin Concentrations, Bioresearch Open Acc 1(1) (2012) 34–40.

[21] O.V. Kim, Z.L. Xu, E.D. Rosen, M.S. Alber, Fibrin Networks Regulate Protein Transport during Thrombus Development, Plos Comput Biol 9(6) (2013) e1003095, 1-10.

[22] S. Muenster, B. Fabry, A Simplified Implementation of the Bubble Analysis of Biopolymer Network Pores, Biophysical Journal 104(12) (2013) 2774–2775.

[23] J. Jesty, The Kinetics of Inhibition of Thrombin by Antithrombin in the Presence of Components of the Hemostatic System, Blood 66(5) (1985) 1189–1195.

[24] W. Li, J. Sigley, S.R. Baker, C.C. Helms, M.T. Kinney, M. Pieters, P.H. Brubaker, R. Cubcciotti, M. Guthold, Nonuniform Internal Structure of Fibrin Fibers: Protein Density and Bond Density Strongly Decrease with Increasing Diameter, Biomed Res Int 2017 (2017) 6385628, 1–13.

[25] S.D. Lewis, P.P. Shields, J.A. Shafer, Characterization of the Kinetic Pathway for Liberation of Fibrinopeptides during Assembly of Fibrin, J Biol Chem 260(18) (1985) 192–199.

[26] M.E. Carr, S.L. Carr, Fibrin Structure and Concentration Alter Clot Elastic-Modulus but Do Not Alter Platelet-Mediated Force Development, Blood Coagulation & Fibrinolysis 6(1) (1995) 79–86.

[27] G. Dietler, W. Kanzig, A. Haeberli, P.W. Straub, Temperature-Dependence of Fibrin Polymerization - a Light-Scattering Study, Biochemistry-Us 24(23) (1985) 6701–6706.

