## supplemental file for "Nonlinear Relationships of Fibrin Network Structure as a Function of Fibrinogen and Thrombin Concentrations for Purified Fibrinogen and Plasma Clots"

#### Table of Contents

**S1. Power Law Justification**

**S2. Structural Median or Mean Data for each Fibrinogen and Thrombin Concentration**

**S3. Additional Analysis to Test Robustness of Statistics (Leave-One-Condition-Out Analysis)**

**S4. Turbidity Analysis of Fibrin Networks Formation with Different Amounts of Added Alexa-488-Labeled Fibrinogen**

### S1. Power Law Justification

#### S1.1. Michaelis–Menten fibrinogen to fibrin conversion with fibrin depletion

We model thrombin cleavage of fibrinogen and subsequent incorporation of fibrin species into growing structures. Let

$F(t) \equiv [\mathrm{Fgn}](t)$: fibrinogen concentration (substrate)

$A(t)$: effective pool of incorporation-competent fibrin species produced by cleavage (fibrin monomers/small oligomers available for incorporation)

$N(t)$: density of reactive growth interfaces (effective growth units/branch sites; a proxy for total reactive interfaces available for incorporation)

$[\mathrm{Thr}]_0$: initial thrombin concentration

$[\mathrm{Thr}]_{\mathrm{eff}}(t) = [\mathrm{Thr}]_0\, a(t)$: effective thrombin level, with $0 < a(t) \leq 1$ capturing inhibition/binding/sequestration without implying stoichiometric thrombin consumption [1]

$k_{\mathrm{cat}}$ and $K_m$: Michaelis–Menten parameters for thrombin cleavage of fibrinogen

$k_{\mathrm{el}}$: effective incorporation (elongation) rate constant

$k_{\mathrm{nuc}}$: effective rate constant for creation of new growth interfaces

$n$: effective exponent summarizing the dominant pathway(s) for interface creation (a coarse-grained, not necessarily integer "order")

$t_*$: fixed readout time

$t_\mathrm{w}$: onset time of the main structure-building window (defined phenomenologically below)

A minimal Michaelis–Menten description of the production rate is

$$v_{\mathrm{prod}}(t) = \frac{k_{\mathrm{cat}}\,[\mathrm{Thr}]_{\mathrm{eff}}(t)\,F(t)}{K_m + F(t)}. \qquad \text{(S1)}$$

Substrate depletion is retained explicitly:

$$\frac{dF}{dt} = -v_{\mathrm{prod}}(t). \qquad \text{(S2)}$$

#### S1.2. Mass balance for $A(t)$ and mass-action incorporation

Fibrin species are produced and incorporated into growing structures:

$$\frac{dA}{dt} = v_{\mathrm{prod}}(t) - v_{\mathrm{cons}}(t). \qquad \text{(S3)}$$

We use a minimal mass-action [2] form for incorporation:

$$v_{\mathrm{cons}}(t) = k_{\mathrm{el}}\,A(t)\,N(t). \qquad \text{(S4)}$$

#### S1.3. Effective creation of growth interfaces

We coarse-grain the creation of new reactive interfaces as

$$\frac{dN}{dt} = k_{\text{nuc}}\, A(t)^{n}, \qquad \text{(S5)}$$

where $n$ summarizes the dominant pathway(s) generating interfaces in an effective description [3].

#### S1.4. Main structure-building window and quasi-steady clamping of $A(t)$

The early-time kinetics of fibrin generation and incorporation are complex and not fully resolved. We therefore introduce a phenomenological main structure-building window $[t_w, t_*]$, defined as the interval that contributes most strongly to the structural readouts at $t_*$. Quasi-steady-state (QSSA) reductions are widely used in enzyme/biochemical kinetics to eliminate fast-relaxing intermediate species [4]; here we use this idea as an effective approximation to remove $A(t)$ from the dynamics and obtain analytic power-law scalings. Specifically, during $[t_w, t_*]$ we assume production and incorporation are of comparable magnitude, so the activated pool $A(t)$ is approximately clamped and satisfies a production–consumption balance:

$$\frac{dA}{dt} \approx 0 \Rightarrow A(t) \approx \frac{v_{\text{prod}}(t)}{k_{\text{el}}\, N(t)}, t \in [t_{\text{w}}, t_*]. \qquad \text{(S6)}$$

Substituting Eq. (S6) into Eq. (S5) gives, for $t \in [t_{\text{w}}, t_*]$,

$$\frac{dN}{dt} = k_{\text{nuc}} \left(\frac{v_{\text{prod}}(t)}{k_{\text{el}}\, N(t)}\right)^{n} \Rightarrow N(t)^{n}\, \frac{dN}{dt} = \frac{k_{\text{nuc}}}{k_{\text{el}}^{n}}\, v_{\text{prod}}(t)^{n}. \qquad \text{(S7)}$$

Integrating over the main window yields

$$N(t_*)^{n+1} - N(t_\mathrm{w})^{n+1} \propto \int_{t_\mathrm{w}}^{t_*} v_\mathrm{prod}\,(t)^n\,dt. \qquad \text{(S8)}$$

The term $N(t_\mathrm{w})^{n+1}$ is an initial-condition contribution at the onset of the window. In practice it can be negligible relative to $N(t_*)^{n+1}$ or absorbed into the prefactor without affecting the scaling exponents; thus, we write the dominant scaling as

$$N(t_*)^{n+1} \propto \int_{t_\mathrm{w}}^{t_*} v_\mathrm{prod}\,(t)^n\,dt. \qquad \text{(S8')}$$

Using $[\mathrm{Thr}]_\mathrm{eff}(t) = [\mathrm{Thr}]_0\,a(t)$ in Eq. (S1),

$$v_\mathrm{prod}(t)^n = (k_\mathrm{cat}[\mathrm{Thr}]_0)^n\,a(t)^n\left(\frac{F(t)}{K_m + F(t)}\right)^n. \qquad \text{(S9)}$$

Therefore,

$$N(t_*)^{n+1} \propto [\mathrm{Thr}]_0^n\,I(F_0; K_m, t_\mathrm{w}, t_*), \qquad \text{(S10)}$$

where the window functional is

$$I(F_0; K_m, t_{\mathrm{w}}, t_*) \equiv \int_{t_{\mathrm{w}}}^{t_*} a(t)^n \left(\frac{F(t)}{K_m + F(t)}\right)^n dt. \qquad \text{(S11)}$$

#### S1.5. Relation between $K_m$ and our concentration range; linear-Michaelis-Menten regime during the main window

In our experiments, the initial fibrinogen concentrations are 0.36–1.45 mg/mL for purified and 0.36–2.9 mg/mL for plasma. Using $M_W(\mathrm{Fgn}) \approx 340$ kDa, these correspond to

$$0.36, 0.73, 1.45, 2.9\, mg/mL \;\Rightarrow 1.06, 2.15, 4.26, 8.53\, \mu M. \qquad \text{(S12)}$$

Using previous reported $K_m$ (a few $\mu$M ) [5], the initial substrate level $F_0$ can be comparable to $K_m$ at $t = 0$. However, because substrate is depleted (Eq. S2), and because the dominant structural contribution is assumed to arise after an initial transient once the system has entered the main window $[t_{\mathrm{w}}, t_*]$ (Eq. S6), so that the substrate level sampled during this window lies in the low-substrate regime:

$$F(t) \ll K_m, t \in [t_{\mathrm{w}}, t_*], \qquad \text{(S13)}$$

then the Michaelis–Menten factor is well approximated by its linear form

$$\frac{F(t)}{K_m + F(t)} \approx \frac{F(t)}{K_m}. \qquad \text{(S14)}$$

Under this linearization, the depletion dynamics are effectively linear in $F$ over the window, so different initial conditions approximately rescale the substrate time course:

$$F(t) = F_0\, g(t), t \in [t_{\mathrm{w}}, t_*], \qquad \text{(S15)}$$

where $g(t)$ represents a shared temporal shape (set by $[\mathrm{Thr}]_0$ and $a(t)$) that is approximately independent of $F_0$ over the finite range studied.

Substituting Eqs. (S14)–(S15) into Eq. (S11) yields

$$I(F_0; K_m, t_{\mathrm{w}}, t_*) \approx \left(\frac{F_0}{K_m}\right)^n \int_{t_{\mathrm{w}}}^{t_*} a(t)^n\, g(t)^n\, dt \equiv \left(\frac{F_0}{K_m}\right)^n J([\mathrm{Thr}]_0; t_{\mathrm{w}}, t_*), \qquad \text{(S16)}$$

with

$$J([\mathrm{Thr}]_0; t_{\mathrm{w}}, t_*) \equiv \int_{t_{\mathrm{w}}}^{t_*} a(t)^n g(t)^n dt. \qquad \text{(S17)}$$

Combining Eqs. (S10) and (S16) gives the scaling

$$N(t_*) \propto [\mathrm{Thr}]_0^{\frac{n}{n+1}}\, F_0^{\frac{n}{n+1}}. \qquad \text{(S18)}$$

**S1.6. Geometric allocation and emergence of the final power law (example: diameter)**

At readout time $t_*$ in a fixed observation volume, the total incorporated fibrin mass scales to leading order with the initial fibrinogen supply:

$$M \propto F_0. \qquad \text{(S19)}$$

Assuming uniform density and equal mass allocation across $N(t_*)$ growth units,

$$M \propto N(t_*)\, d^2\, \ell(t_*), \qquad \text{(S20)}$$

where $d$ is fiber diameter and $\ell(t_*)$ is an effective length contributed per growth unit at readout. If $\ell$ varies weakly across the matrix or follows an effective power law

$$\ell(t_*) \propto F_0^u\, [\mathrm{Thr}]_0^v, \qquad \text{(S21)}$$

then combining Eqs. (S18)–(S21) yields the multiplicative power law

$$d \propto F_0^\alpha\, [\mathrm{Thr}]_0^\beta, \qquad \text{(S22)}$$

with

$$\alpha = \frac{1}{2}\left(1 - \frac{n}{n+1} - u\right) = \frac{1}{2}\left(\frac{1}{n+1} - u\right), \qquad \text{(S23)}$$

$$\beta = -\frac{1}{2}\left(\frac{n}{n+1} + v\right). \qquad \text{(S24)}$$

#### S2. Structural Median or Mean Data for each Fibrinogen and Thrombin Concentration

##### S2.1. Fiber Diameter

| Fgn (mg/mL) | Thr (U/mL) | Plasma Fiber Diameter Median [IQR] (n=16) | Purified Fiber Diameter Median [IQR] (n=16) |
|---|---|---|---|
| 0.36 | 0.05 | 130 [127, 136] | 100 [98, 107] |
| 0.36 | 0.1 | 128 [114, 138] | 91 [89, 93] |
| 0.36 | 0.2 | 100 [92, 113] | 76 [73, 78] |
| 0.36 | 1 | 61 [59, 67] | |
| 0.73 | 0.05 | 142 [140, 151] | 116 [110, 119] |
| 0.73 | 0.1 | 131 [126, 138] | 102 [99, 108] |
| 0.73 | 0.2 | 117 [111, 127] | 83 [81, 87] |
| 0.73 | 1 | 72 [69, 77] | |
| 1.45 | 0.05 | 160 [150, 171] | 125 [117, 127] |
| 1.45 | 0.1 | 159 [147, 167] | 115 [107, 121] |
| 1.45 | 0.2 | 130 [125, 136] | 93 [92, 96] |
| 1.45 | 1 | 91 [87, 93] | |
| 2.9 | 0.05 | 190 [177, 195] | |
| 2.9 | 0.1 | 162 [144, 186] | |
| 2.9 | 0.2 | 154 [145, 168] | |
| 2.9 | 1 | 98 [93, 115] | |

*Table S1:* Median fiber diameters for plasma and purified fibrin networks, calculated from per-image medians across 16 analyzed images; interquartile ranges (IQR) are reported for each condition.

**S2.2. Fiber Length**

| Fgn (mg/mL) | Thr (U/mL) | Plasma Fiber Length Median [IQR] (n=16) | Purified Fiber Length Median [IQR] (n=16) |
|---|---|---|---|
| 0.36 | 0.05 | 2143 [1942, 2414] | 1686 [1608, 1811] |
| 0.36 | 0.1 | 1750 [1567, 1940] | 1328 [1210, 1483] |
| 0.36 | 0.2 | 1624 [1455, 1969] | 1114 [974, 1216] |
| 0.36 | 1 | 1093 [1016, 1216] | |
| 0.73 | 0.05 | 2302 [1836, 2726] | 1786 [1617, 1910] |
| 0.73 | 0.1 | 1848 [1694, 2189] | 1507 [1285, 1605] |
| 0.73 | 0.2 | 1554 [1430, 1726] | 1262 [1123, 1367] |
| 0.73 | 1 | 1087 [995, 1251] | |
| 1.45 | 0.05 | 2379 [2267, 2664] | 1979 [1730, 2029] |
| 1.45 | 0.1 | 1934 [1713, 2267] | 1516 [1399, 1594] |
| 1.45 | 0.2 | 1745 [1571, 1876] | 1259 [1135, 1329] |
| 1.45 | 1 | 1162 [1019, 1299] | |
| 2.9 | 0.05 | 2528 [2173, 2781] | |
| 2.9 | 0.1 | 2119 [2076, 2373] | |
| 2.9 | 0.2 | 1891 [1762, 1968] | |
| 2.9 | 1 | 1297 [1170, 1384] | |

*Table S2:* Median fiber lengths for plasma and purified fibrin networks, calculated from per-image medians across 16 analyzed images; interquartile ranges (IQR) are reported for each condition.

**S2.3. Fiber Density**

| Fgn (mg/mL) | Thr (U/mL) | Plasma Fiber Density Mean ± SD (n=36) | Purified Fiber Density Mean ±SD (n=36) |
|---|---|---|---|
| 0.36 | 0.05 | 5.1 ± 0.8 | 7.3 ± 0.8 |
| 0.36 | 0.1 | 5.9 ± 0.9 | 10.0 ± 1.4 |
| 0.36 | 0.2 | 9.7 ± 1.2 | 15.1 ± 1.4 |
| 0.36 | 1 | 21 ± 2 | |
| 0.73 | 0.05 | 10.2 ± 2.1 | 12.7 ± 1.2 |
| 0.73 | 0.1 | 10.7 ± 0.9 | 16.0 ± 1.5 |
| 0.73 | 0.2 | 11.9 ± 1.2 | 22 ± 2 |
| 0.73 | 1 | 24 ± 2 | |
| 1.45 | 0.05 | 14 ± 2 | 22 ± 2 |
| 1.45 | 0.1 | 15.5 ± 1.6 | 29 ± 2 |
| 1.45 | 0.2 | 14.0 ± 1.6 | 36 ± 2 |
| 1.45 | 1 | 27 ± 3 | |
| 2.9 | 0.05 | 18 ± 2 | |
| 2.9 | 0.1 | 20 ± 5 | |
| 2.9 | 0.2 | 22 ± 4 | |
| 2.9 | 1 | 32 ± 4 | |

*Table S3:* Mean fiber densities for plasma and purified fibrin networks, calculated from per-image means across 36 analyzed images; standard deviation (SD) are reported for each condition.

#### S2.4. Bubble Diameter

| Fgn (mg/mL) | Thr (U/mL) | Plasma Bubble Diameter Median [IQR] (n=36) | Purified Bubble Diameter Median [IQR] (n=36) |
|---|---|---|---|
| 0.36 | 0.05 | 13.7 [12.9, 13.9] | 12.0 [11.4, 12.5] |
| 0.36 | 0.1 | 13.1 [11.4, 13.6] | 11.1 [10.8, 11.5] |
| 0.36 | 0.2 | 10.5 [10.2, 11.2] | 8.9 [8.6, 9.3] |
| 0.36 | 1 | 7.0 [6.8, 7.2] | |
| 0.73 | 0.05 | 10.1 [10.0, 10.3] | 8.5 [8.2, 8.7] |
| 0.73 | 0.1 | 9.9 [9.6, 10.3] | 8.2 [7.7, 8.4] |
| 0.73 | 0.2 | 9.3 [8.6, 9.7] | 7.6 [7.3, 7.8] |
| 0.73 | 1 | 6.7 [6.6, 6.9] | |
| 1.45 | 0.05 | 8.6 [8.0, 9.0] | 6.6 [6.4, 6.8] |
| 1.45 | 0.1 | 7.7 [7.0, 8.0] | 5.9 [5.8, 6.0] |
| 1.45 | 0.2 | 6.9 [6.3, 7.0] | 5.60 [5.56, 5.64] |
| 1.45 | 1 | 6.4 [6.2, 6.5] | |
| 2.9 | 0.05 | 7.2 [6.6, 8.1] | |
| 2.9 | 0.1 | 6.9 [6.5, 7.2] | |
| 2.9 | 0.2 | 6.6 [6.4, 6.9] | |
| 2.9 | 1 | 5.9 [5.7, 6.1] | |

*Table S4:* Median bubble diameter for plasma and purified fibrin networks, calculated from per-image medians across 36 analyzed images; interquartile ranges (IQR) are reported for each condition.

**S3. Additional Analysis to Test Robustness of Statistics (Leave-One-Condition-Out Analysis)**

| Plasma | | Value | Leave-One-Condition-Out |
|---|---|---|---|
| Parameters for Fiber diameter | k | 81 | [79, 84] |
| | α | 0.19 | [0.17, 0.20] |
| | β | - 0.23 | [-0.25, -0.21] |
| Parameters for Fiber length | k | 1155 | [1144, 1166] |
| | α | 0.09 | [0.08, 0.09] |
| | β | -0.24 | [-0.24, -0.23] |
| Parameters for Fiber density | k | 25 | [22, 25] |
| | α | 0.45 | [0.40, 0.48] |
| | β | 0.30 | [0.26, 0.32] |
| Parameters for Bubble diameter | k | 6.6 | [6.5, 6.6] |
| | α | -0.24 | [-0.25, -0.22] |
| | β | -0.14 | [-0.15, -0.13] |

*Table S5:* Power-law fitting parameters and leave-one-condition-out ranges for plasma clots

| Purified Fibrinogen | | Value | Leave-One-Condition-Out |
|---|---|---|---|
| Parameters for Fiber diameter | k | 64 | [61, 65] |
| | α | 0.15 | [0.15, 0.17] |
| | β | - 0.21 | [-0.23, -0.21] |
| Parameters for Fiber length | k | 775 | [741, 797] |
| | α | 0.07 | [0.07, 0.11] |
| | β | -0.29 | [-0.31, -0.28] |
| Parameters for Fiber density | k | 57 | [51, 59] |
| | α | 0.73 | [0.69, 0.75] |
| | β | 0.43 | [0.38, 0.44] |
| Parameters for Bubble diameter | k | 5.1 | [4.9, 5.5] |
| | α | -0.41 | [-0.44, -0.39] |
| | β | -0.14 | [-0.16, -0.11] |

*Table S6:* Power-law fitting parameters and leave-one-condition-out ranges for purified fibrinogen clots

#### S4. Turbidity Analysis of Fibrin Networks Formation with Different Amounts of Added Alexa-488-Labeled Fibrinogen

**Methods.** Turbidity-based clot formation assays were performed using a SpectraMax microplate reader by monitoring absorbance at 405 nm every 7 s for 3 h in 96-well microplates (Corning flat clear-bottom white polystyrene, TC-treated). Plasma clots were prepared using fibrinogen concentrations of 0.73, 1.45, or 2.4 mg/mL with thrombin fixed at 0.2 U/mL. To evaluate the effect of fluorescent labeling on clot formation, Alexa Fluor 488-labeled fibrinogen was incorporated at 0%, 1.5%, 3%, 6%, or 12% of the total fibrinogen concentration.

Each curve is the average of two separate experiments.

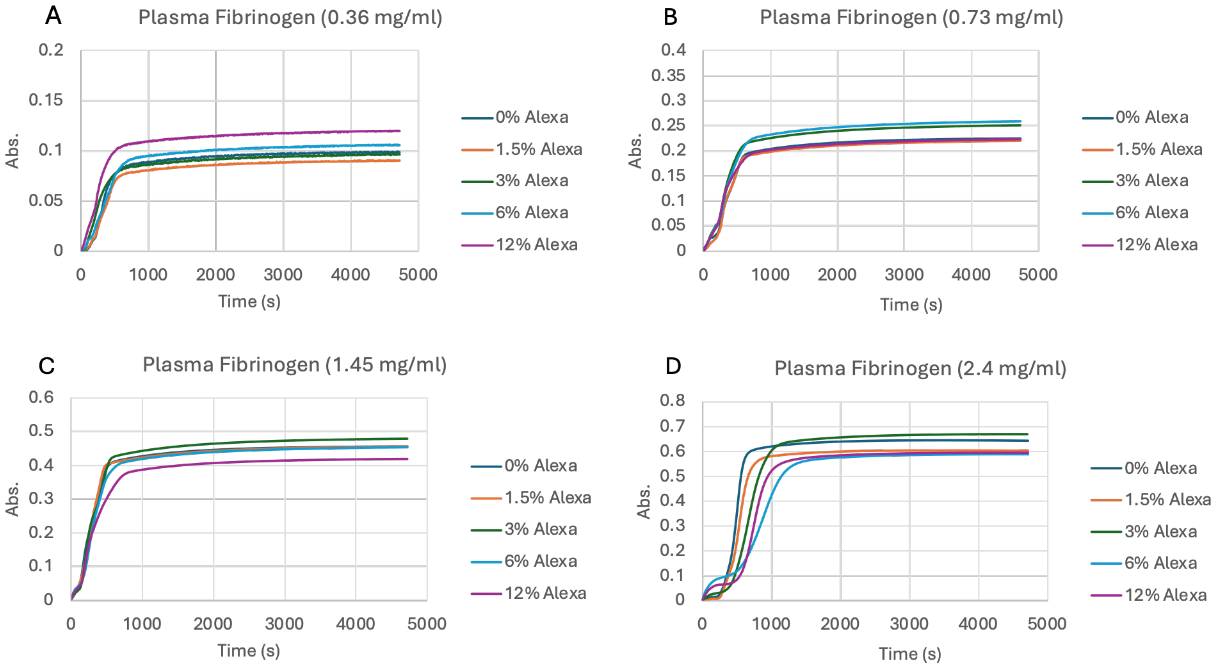


*Figure S1. Turbidity curves for plasma fibrinogen concentrations ranging from 0.36 to 2.4 mg/mL with 0–12% Alexa Fluor 488 labeled fibrinogen added.*

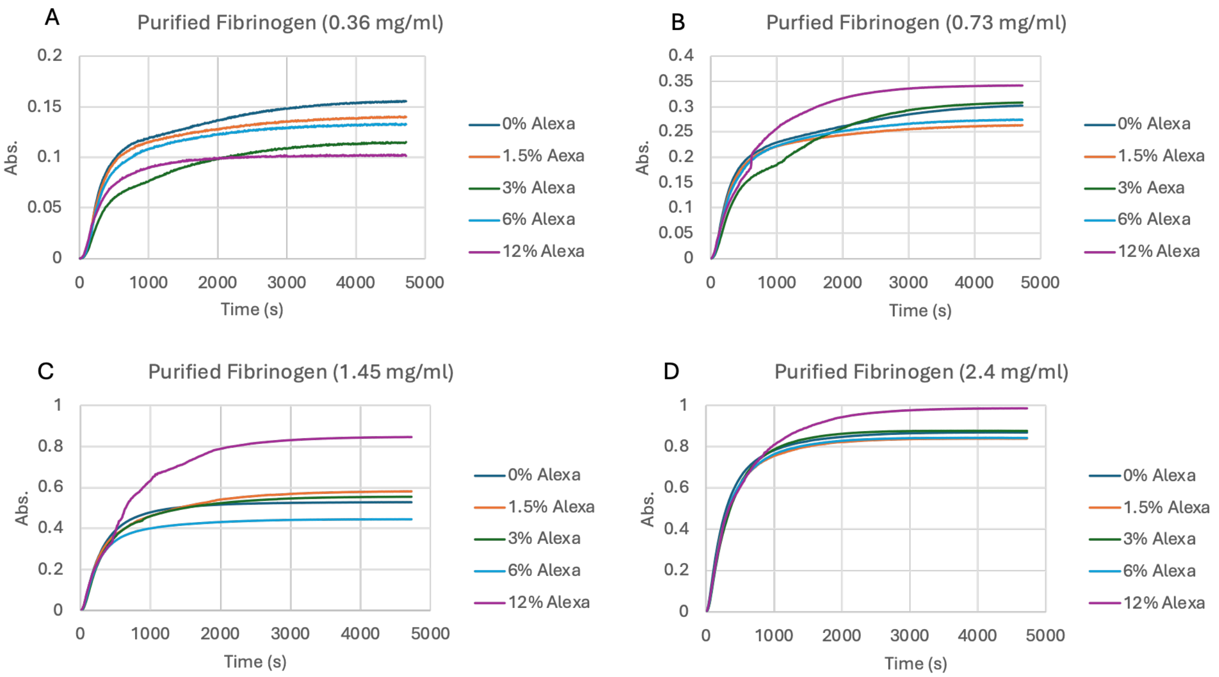


*Figure S2. Turbidity curves for purified fibrinogen concentrations ranging from 0.36 to 2.4 mg/mL with 0–12% Alexa Fluor 488 labeled fibrinogen added.*

Minimal differences in fibrin formation dynamics and final turbidity were observed up to 6% labeling, whereas more noticeable deviations appeared at 12% labeling.

#### References

[1] J. Jesty, The Kinetics of Inhibition of Thrombin by Antithrombin in the Presence of Components of the Hemostatic System, Blood 66(5) (1985) 1189-1195.

[2] J.W. Weisel, C. Nagaswami, Computer Modeling of Fibrin Polymerization Kinetics Correlated with Electron-Microscope and Turbidity Observations - Clot Structure and Assembly Are Kinetically Controlled, Biophysical Journal 63(1) (1992) 111-128.

[3] A.L. Fogelson, J.P. Keener, Toward an understanding of fibrin branching structure, Phys Rev E 81(5) (2010) 051922, 1-9.

[4] L.A. Segel, M. Slemrod, The Quasi-Steady-State Assumption - a Case-Study in Perturbation, Siam Rev 31(3) (1989) 446-477.

[5] D.L. Higgins, S.D. Lewis, J.A. Shafer, Steady-State Kinetic-Parameters for the Thrombin-Catalyzed Conversion of Human-Fibrinogen to Fibrin, J Biol Chem 258(15) (1983) 9276-9282.
